# Mucin-type *O*-glycans regulate proteoglycan stability and chondrocyte maturation

**DOI:** 10.64898/2025.12.11.693745

**Authors:** Xiaolin Dong, Sydney Bedillion, Kaleigh E. Gosnell, Peng Zhao, Amrita Basu, Kavya Suryadevara, Digantkumar Chapla, Lance Wells, Ryan J. Weiss

## Abstract

*O*-glycosylation is a ubiquitous post-translational modification essential for protein stability, cell signaling, and tissue organization, yet how distinct *O*-glycan subclasses coordinate tissue development remains unclear. Here, we identify functional crosstalk between extended mucin-type *O*-glycans and heparan sulfate proteoglycans (HSPGs). Genetic ablation of β-1,3-galactosyltransferase 1 (C1GALT1) or its chaperone COSMC in human chondrocytes reduced cell surface HSPGs and fibroblast growth factor (FGF) binding, leading to impaired MAPK/ERK signaling, which was recapitulated in hypomorphic *COSMC-CDG* patient cells. Transcriptomic and secretome analyses revealed selective loss of proteoglycan expression and broader extracellular matrix remodeling in COSMC*-* and C1GALT1-deficient cells. Mechanistically, truncation of *O*-GalNAc glycans reduced Syndecan-1 levels at the cell surface via enhancing its lysosome-dependent degradation and diminished CD44v3-mediated FGF1 binding. Functionally, loss of complex *O*-GalNAc glycans disrupted chondrogenesis of growth plate-like chondroprogenitors. Overall, these findings reveal previously unrecognized roles for mucin-type *O*-glycans in maintaining proteoglycan stability and function, highlighting cross-regulatory mechanisms of glycosylation crucial for normal development that likely contribute to the pathophysiology of glycosylation-related disorders.

## Introduction

Glycosylation confers structural complexity to the proteome, directing protein folding, trafficking, and stability while orchestrating processes essential for tissue development and homeostasis^1, 2^. Increasing evidence indicates that distinct glycosylation pathways are interconnected, with one type of glycan influencing the efficiency, content, and spatial organization of others^3^. This coordination could arise through shared nucleotide-sugar precursors and/or diverse forms of glycosylation on the same protein, which together can influence protein assembly, trafficking, and function. Such dynamic cross-regulation among glycans is increasingly recognized as a key determinant of cell surface organization, receptor recognition, and cellular function^4, 5^, yet the molecular mechanisms governing this interplay remain poorly understood.

Among the diverse forms of glycosylation, *O*-glycans represent the most common covalent modifications of serine and threonine residues of mammalian glycoproteins. Two major subclasses of *O*-glycans, mucin type *O*-GalNAc glycans and glycosaminoglycans (GAGs), exhibit distinct but complementary roles in tuning protein function and extracellular matrix (ECM) assembly^5^. *O*-GalNAc glycans are among the most abundant and diverse types of *O*-glycosylation. *O*-GalNAcylation is initiated in the Golgi apparatus by polypeptide GalNAc-transferases (ppGalNAc-Ts) enzymes. These enzymes attach *N*-acetylgalactosamine (GalNAc) to serine or threonine residues, forming the Tn antigen. Next, the core 1 β1,3-galactosyltransferase (C1GALT1), along with its obligate molecular chaperone, COSMC (*C1GALT1C1*, an X-linked gene on Xq24), adds galactose to form the T antigen (core 1 *O*-glycan). This core 1 structure serves as the required platform for the generation of a large assortment of elaborated core structures to produce the full array of mature *O*-glycans^6^. Mature mucin-type O-glycans regulate protein folding, secretion, protein-glycan binding, and cell-cell interactions, whereas their truncation is a hallmark of several diseases, including cancer and rare genetic disorders^7, 8^. Importantly, *O*-GalNAc glycans can also act as Golgi export signals, which influences the secretion, stability, and localization of glycoproteins and tissue organization^9^.

Glycosaminoglycans (GAG) are long, linear polysaccharides consisting of repeating glucosamine (GlcN) and glucuronic/iduronic acid (GlcA/IdoA) subunits that are variably sulfated and *O*-linked to serine residues of core proteins, known as proteoglycans (PGs)^10, 11^. Two principal GAG subclasses, heparan sulfate (HS) and chondroitin/dermatan sulfate (CS/DS), are essential components of the cell surface and extracellular matrix (ECM) and are ubiquitously expressed in animal tissues. HS and CS/DS proteoglycans collectively regulate cell signaling and homeostasis by binding growth factors, integrins, and ECM proteins^12^. Their structural diversity in chain length and sulfation pattern fine-tunes these interactions, controlling cell adhesion, migration, and tissue morphogenesis^10^. In skeletal tissues, for example, proteoglycans form complexes with collagen and hyaluronan that provide structural integrity and guide cartilage and bone development^13^.

Defects in *O*-GalNAc or GAG biosynthesis underlie a subset of rare genetic disorders, known as congenital disorders of glycosylation (CDGs), which manifest with pleiotropic effects on cell function across tissues^14, 15^. Many CDG disorders exhibit pleiotropic overlapping clinical features, including developmental disabilities, multi-organ involvement, short stature, and skeletal abnormalities^16^. For example, a recent study revealed that *COSMC*-CDG patients, caused by inherited mutations in *COSMC/C1GALT1C1 (*A20D-COSMC), frequently exhibit skeletal defects and short stature in addition to other systemic issues^15^. Similarly, *EXT1/EXT2*-CDG, commonly known as Multiple Hereditary Exostoses (MHE), results from heterozygous mutations in *EXT1* or *EXT2*, which encode glycosyltransferases essential for heparan sulfate elongation, leading to the formation of bony outgrowths that cause chronic pain and can interfere with normal bone development^17^. Interestingly, a recent study reported that mice with chondrocyte-specific transgenic expression of *Galnt3*, a polypeptide GalNAc transferase encoding gene, exhibit enhanced Tn antigen levels at the growth plate, which coincided with decreased GAG levels, dwarfism, reduced chondrocyte maturation, and skeletal dysplasia^18^. These findings point to functional cross-regulation between *O*-GalNAc glycosylation and proteoglycan assembly. Additionally, the convergence of skeletal defects across distinct CDGs implies a shared requirement for coordinated *O*-glycan and proteoglycan regulation during development.

Although the majority of membrane-bound and secreted proteoglycans also carry *O*-GalNAc glycans^19^, direct mechanistic evidence linking *O*-GalNAc glycosylation to proteoglycan stability and function remains lacking. Here, we tested the hypothesis that mucin-type O-glycans regulate proteoglycan trafficking and cell surface presentation. Using CRISPR/Cas9-mediated knockout of *COSMC* or *C1GALT1*, respectively, in human and murine chondrocytes, we demonstrate that truncation of mucin-type *O*-GalNAc glycans destabilizes cell-surface proteoglycans and alters ECM composition. Transcriptomic and secretomic profiling indicated global remodeling of proteoglycan biosynthesis and secretion in COSMC/C1GALT1 deficient cells, while in vitro differentiation assays using a cell-based model of the growth plate revealed impaired chondrogenic maturation. Together, these findings uncover a fundamental cross-regulatory mechanism linking mucin-type *O*-glycosylation and proteoglycan function that reveals new insights into *O*-glycan functions in development and glycosylation-related disorders.

## Results

### Global truncation of *O*-GalNAc glycans reduces HSPGs and impairs HS-dependent growth factor binding and signaling

To assess whether mucin-type *O*-GalNAc glycosylation influences proteoglycan biosynthesis and cell surface presentation, we genetically disrupted the core 1 *O*-glycan biosynthetic pathway in human chondrocytes. For our studies, we utilized immortalized TC28a2 human chondrocytes, which highly express cell-surface proteoglycans and serve as a tractable in vitro model to study the cross regulation of *O*-GalNAc glycans and proteoglycans in cartilage^20,21^. With over 20 known ppGalNAc-Ts (GALNTs), genetic redundancy makes disrupting *O*-GalNAc initiation via a single gene knockout challenging^22, 23^. In addition, studies have shown that loss of individual GALNTs often leads to substrate-specific effects rather than global abolition of *O*-GalNAcylation^24^. Instead, we utilized CRISPR/Cas9 to globally truncate *O*-GalNAc glycans through genetic ablation of β1,3-galactosyltransferase (*C1GALT1*) or its chaperone, COSMC (*C1GALT1C1*), which coordinate to add galactose (Gal) to GalNAc to generate the T antigen at serine/threonine residues of the core protein and thus eliminates the production of both Core 1 and Core 2 extended *O*-GalNAc glycans^6^ (Fig. 1A). Successful gene disruption was confirmed by Western blotting and Sanger sequencing (Fig. 1B-C, Suppl. Fig. 1A-B). Flow cytometry following pre-treatment with sialidase demonstrated robust accumulation of the Tn antigen in both knockout lines, detected by VVA lectin and anti-Tn antibody cell surface staining, (Fig. 1D; Suppl. Fig. 1C), accompanied by the expected corresponding loss of the Core 1 T antigen detected by PNA lectin binding (Fig. 1E).

**Figure 1.**
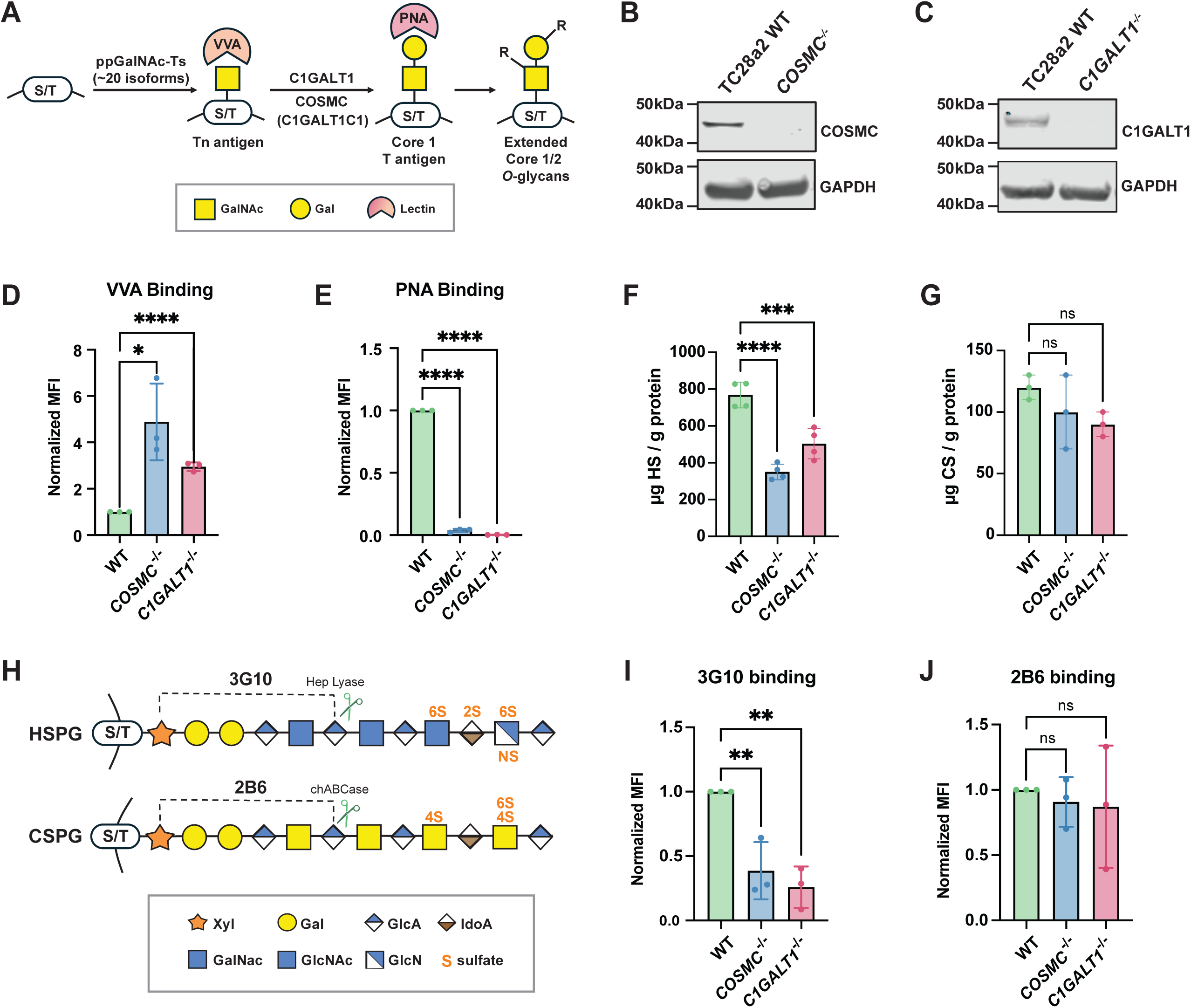
Genetic ablation of *COSMC/C1GALT1* reduces cell surface heparan sulfate in TC28a2 chondrocytes. (A) Schematic of mucin-type *O*-GalNAcylation. Polypeptide GalNAc-transferases (ppGalNAc-Ts/GALNTs; ∼20 isoenzymes) initiate addition of GalNAc to serine or threonine residues of core proteins. Loss of COSMC or C1GALT1 blocks core 1 extension (T antigen), resulting in accumulation of truncated *O*-glycans (Tn antigen). VVA lectin detects Tn antigen, whereas PNA lectin detects T antigen. Western blot validation of (B) *COSMC^-/-^*and (C) *C1GALT1^-/-^* knockout cell lines; GAPDH serves as a loading control. (D-E) Flow cytometry analysis of VVA and PNA lectin to wild-type and COSMC/C1GALT1 knockout cells. (F-G) LC-MS quantification of total cell surface heparan sulfate (HS) and chondroitin sulfate (CS) levels. (H) Schematic of GAG-specific antibodies: 3G10 recognizes HS stubs generated by heparin lyase and 2B6 detects CS stubs generated by chondroitinase ABC. (I) Flow cytometry analysis of 3G10 binding in wild-type and knockout cells. (J) Flow cytometry analysis of 2B6 binding in wild-type and knockout cells. Data points are shown as mean ± SD (n ≥ 3 independent experiments); p-values were determined by one-way ANOVA with Tukey post-test with ****p < 0.0001, *** p < 0.001, ** p < 0.01, * p < 0.05.

After confirming truncation of extended *O*-GalNAc glycans, we purified cell surface HS and CS/DS polysaccharides from wild-type, *COSMC*^-/-^, and *C1GALT1*^-/-^ cells via anion exchange chromatography and analyzed their disaccharide composition using liquid chromatography coupled to mass spectrometry (LC-MS), using previously reported methods (Suppl. Fig 1D)^25, 26^. Interestingly, these analyses revealed a reduction in total HS levels in both *COSMC*^-/-^ and *C1GALT1*^-/-^ knockout cells with no significant change in CS/DS content (Fig. 1F-G). HS disaccharide composition remained largely unchanged, with a slight reduction in 6-*O* sulfated disaccharides in the *COSMC^-/-^* knockout line (Suppl. Fig 1E-G), while CS/DS disaccharides were largely unchanged in both knockout lines, with a slight decrease in 6-O sulfation in *C1GALT1*^-/-^cells (Suppl. Fig. 1H-I). To clarify these findings, we conducted flow cytometry using proteoglycan-specific antibodies, including 3G10, which recognizes HS neoepitopes on proteoglycans produced upon heparinase digestion^27^, and 2B6, which detects CS neoepitopes after chondroitinase ABC digestion^28^ (Fig. 1H). Flow cytometry analysis revealed a ∼80% reduction in 3G10 binding with no significant changes in 2B6 binding (Fig. 1I-J), indicating that the observed decreased HS levels were due to a reduction in HSPGs and/or HS-modified sites at the cell surface. Overall, these findings reveal that truncation of *O*-GalNAc glycans results in a selective reduction in HSPG presentation at the cell surface in human chondrocytes.

HSPGs are known to function as ternary receptors for fibroblast growth factors (FGFs) and their fibroblast growth factor receptors (FGFRs), thus aiding in receptor-mediated activation of downstream signaling pathways, including MAPK, relevant for musculoskeletal development (Fig. 2A)^29, 30^. Given that we observed a selective decrease in cell surface HS in *COSMC*^-/-^ and *C1GALT1*^-/-^ cells, we hypothesized that truncation of *O*-GalNAc glycans would impair HS-dependent FGF binding and downstream MAPK signaling. Flow cytometry analysis revealed a significant decrease in cell surface binding of two well-characterized HS-binding proteins, fibroblast growth factor 1 (FGF1) and fibroblast growth factor 2 (FGF2)^12, 31^, in *COSMC*^-/-^ and C*1GALT1*^-/-^ knockout cells (Fig. 2B-C). We independently confirmed these observed FGF binding defects in additional *COSMC*^-/-^ and *C1GALT1*^-/-^ knockout clonal lines (*COSMC*^c6^, *C1GALT1*^t2^), which also exhibited decreased 3G10 binding and increased Tn antigen levels (Suppl. Fig. 2). To examine whether reduced growth factor binding was driven by alterations in FGF receptor (FGFR) expression, we measured FGFR1 cell surface levels, which is highly expressed in TC28a2 cells, via flow cytometry using an FGFR1-specific antibody. This analysis revealed minimal differences in FGFR1 levels between wild-type and knockout lines (Fig. 2D), suggesting an HS-dependent reduction in FGF binding at the cell surface. Consistent with reduced cellular HS levels and FGF binding, FGF2-induced ERK1/2 phosphorylation was also significantly reduced in *COSMC*^-/-^ and *C1GALT1*^-/-^ knockout cells compared to wild-type controls (Fig. 2E-F).

**Figure 2.**
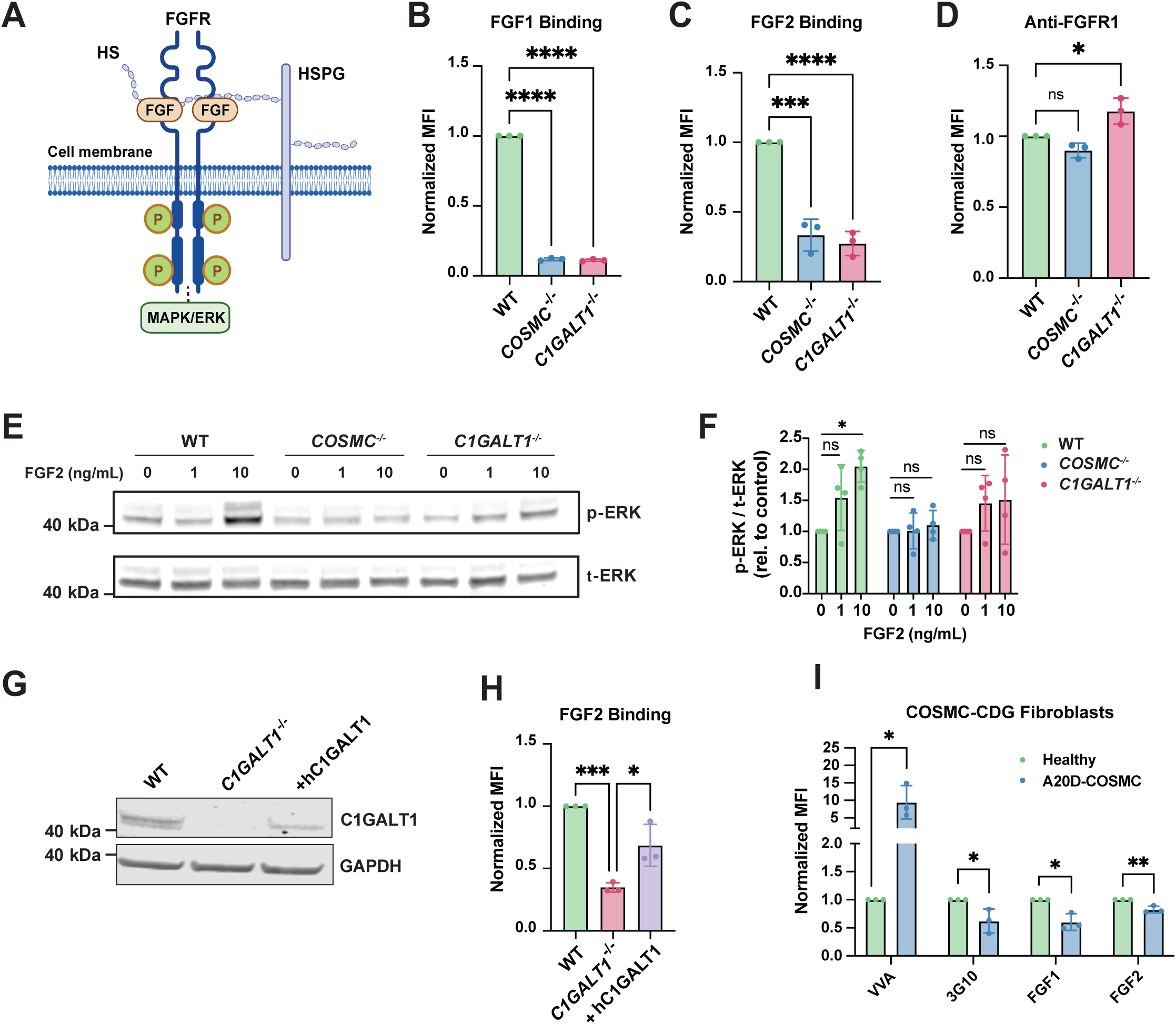
Truncation of *O*-GalNAc glycans disrupts FGF binding and downstream MAPK/ERK signaling. (A) Diagram of the FGF-FGFR-heparan sulfate ternary complex, whch regulates downstream pathways including MAPK/ERK signaling. (B-C) FGF1 and FGF2 ligand binding were significantly reduced in knockout cells versus wild-type control cells. (D) Cell surface expression of FGFR1 was unchanged in wild-type and knockout cells, as quantified via flow cytometry. (E) Western blot analysis of FGF2 induced ERK phosphorylation (p-ERK) and total ERK (t-ERK) levels in wild-type and knockout cells. (F) Western blot band intensities were quantitated by densitometry and plotted as the ratio of p-ERK to t-ERK signal. (G) Western blot analysis of C1GALT1 protein levels upon reintroduction of human C1GALT1 (hC1GALT1) into *C1GALT1^-/-^*cells. (H) Flow cytometry analysis of FGF2 binding in wild-type, *C1GALT1^-/-^*, and *C1GALT1^-/-^* cells transduced with *C1GALT1* cDNA. (I) Flow cytometry analysis of HS stubs using 3G10 antibody, and FGF binding in A20D-*COSMC-CDG* patient fibroblasts. Data points are shown as mean ± SD (n ≥ 3 independent experiments); p-values were determined by one-way ANOVA with Tukey post-test (A-H) and student’s T-tests (I) with ****p < 0.0001, *** p < 0.001, ** p < 0.01, * p < 0.05.

To rule out off-target effects driving the observed phenotypes, full-length human C1GALT1 cDNA was reintroduced in *C1GALT1^-/-^* knockout cells (Fig. 2G), which restored cell surface binding of FGF2 (Fig. 2H), thus linking truncation of *O*-GalNAc glycans to a decrease in cell surface HS content and function. Importantly, we also validated our findings in primary fibroblasts isolated from a male *COSMC-CDG* patient with the hemizygous variant c.59C>A (p.Ala20Asp) in *C1GALT1C1* (A20D-COSMC). These cells were reported to exhibit similar defects in *O*-glycan extension and accumulate Tn antigen^15^, which we confirmed by VVA binding (Fig. 2I). Intriguingly, the *COSMC-CDG* patient-derived cells also exhibited a reduction in 3G10, FGF1, and FGF2 binding, albeit with more subtle defects compared to TC28a2 knockout lines, which we attributed to low residual T-synthase enzyme activity, as previously reported^15^. Collectively, these results establish *O*-GalNAcylation as a key regulator of HS-mediated growth factor binding and signaling, with critical implications for cell growth and signaling.

### Mucin-type *O*-GalNAc glycans regulate proteoglycan expression and secretion

We next aimed to understand the underlying molecular mechanisms leading to reduced proteoglycan cell surface presentation in COSMC/C1GALT1 deficient cells. First, we performed RNA sequencing on wild-type, *COSMC*^-/-^, and *C1GALT1*^-/-^ knockout cells to compare their transcriptome profiles. A heat map of global differential expression changes revealed distinct hierarchical clustering for *COSMC*^-/-^ and *C1GALT1*^-/-^ knockout cells compared to wild-type, with 128 overlapping genes significantly downregulated and 219 genes upregulated (Fold change ≥ 2, FDR p ≤ 0.01) in both knockout cells (Fig. 3A-B, Suppl. Fig. 3A-B). Gene ontology analysis of the shared differentially expressed gene sets revealed downregulation of pathways related to *extracellular matrix organization*, *retrograde signaling*, and *syndecan interactions* as well as upregulation of pathways related to the *inflammatory response*, *tube morphogenesis*, and *transcriptional misregulation*, consistent with broad effects of *O*-glycan truncation on transcription and associated cellular processes (Fig. 3C-D). When we compared specific expression changes for all GAG-related enzymes and proteoglycans, we found that the largest subset of differentially expressed GAG genes (FDR p ≤ 0.01) encoded proteoglycans, including downregulation of several HSPGs (*SDC2, HSPG2, GPC1, GPC6, COL18A1*) and upregulation of “part-time” proteoglycans, *CD44* and *ESM1*^32^, among others (Fig. 3E). In addition, expression of a subset of GAG biosynthetic genes was altered in both knockout lines, including downregulation of the secreted HS endosulfatases, *SULF1/SULF2,* and upregulation of CS biosynthetic enzymes, *CHST11* and *CSGALNACT1* (Fig. 3E). The top differentially expressed hits in this set (*SULF1, SULF2, SDC2, ESM1*) were also confirmed in the additional *COSMC^c^*^6^ and *C1GALT1^t^*^2^ knockout lines via quantitative PCR (Suppl. Fig. 3C). Among the top hits, we focused on one of the most significantly downregulated genes, Syndecan-2 (*SDC2*), which encodes a key transmembrane HSPG that is a known co-receptor for FGFs and plays roles in tissue development and repair^33^. We orthogonally confirmed a decrease in cell surface SDC2 levels in both knockout lines via flow cytometry using an anti-SDC2 antibody (Fig. 3F), consistent with the observed reduction in mRNA levels (Fig. 3E). Notably, we assessed cell surface levels of a related syndecan protein, Syndecan-1 (SDC1), which exhibited a slight increase in mRNA levels with a surprising significant decrease in cell surface levels in *COSMC^-/-^* and *C1GALT1^-/-^* knockout cells (Fig. 3G). This discrepancy between *SDC1* transcript level changes and reduced protein abundance at the cell surface suggested that SDC1 protein may be regulated post-translationally in cells lacking complex *O*-GalNAc glycans, possibly via enhanced shedding and/or reduced cell surface trafficking.

**Figure 3.**
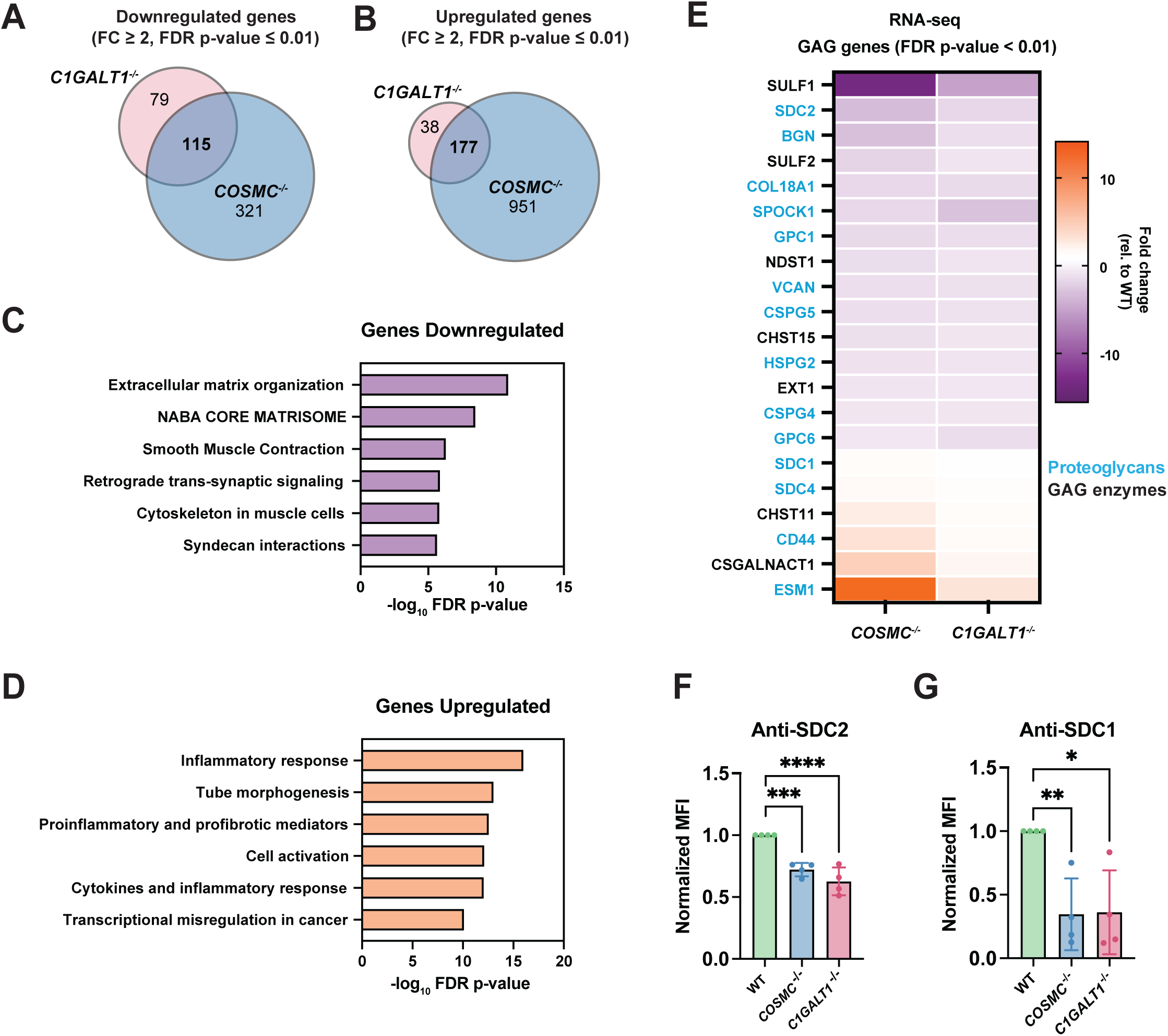
Transcriptome analysis of *COSMC^-/-^* and *C1GALT1^-/-^* knockout cells reveals altered expression of extracellular matrix and proteoglycan genes. Venn diagrams comparing significantly downregulated (A) and upregulated (B) genes (Fold change ≥ 2, FDR p ≤ 0.01) from RNA-seq datasets for *COSMC^-/-^*and *C1GALT1^-/-^* knockout cells. Gene Ontology enrichment analysis of overlapping 128 downregulated (C) and 219 upregulated (D) genes shared in *COSMC^-/-^* and *C1GALT1^-/-^* cells. (E) Differential expression analysis of proteoglycan and GAG biosynthetic genes (FDR p ≤ 0.01) compared to wild-type control cells. Flow cytometry analysis of (F) syndecan-1 (SDC1) and (G) syndecan-2 levels in TC28a2 wild-type, *COSMC^-/-^*, and *C1GALT1^-/-^* cells. Data points are shown as mean ± SD (n ≥ 3 independent experiments); p-values were determined by one-way ANOVA with Tukey post-test with **** p < 0.0001, *** p < 0.001, ** p < 0.01, * p < 0.05.

Prior work has highlighted the importance of *O*-GalNAc glycosylation for the trafficking and secretion of extracellular proteins, which directly influences ECM composition and impacts cell adhesion, growth and organogenesis^34, 35, 36^. Additionally, glycomics-based studies using the SimpleCell technology previously identified *O*-GalNAc modified sites on multiple proteoglycans, including SDC1^23, 37^. To identify the repertoire of *O*-GalNAc modified proteoglycans in TC28a2 chondrocytes, we adapted our selective exoenzymatic labeling (SEEL) method^38^ to detect Tn antigen modified sites in the *COSMC/C1GALT1* knockout lines using CMP-sialic acid-biotin (CMP-sia-biotin) and recombinant ST6GALNAC1 (Suppl. Fig. 4A), which adds a sialic acid to *O*-GalNAc of the Tn antigen^39^. Cells were incubated with sialidase and labeled with ST6GALNAC1 in the presence of CMP-sia-biotin followed by streptavidin enrichment (Suppl. Fig. 4B). Both knockout lines gave substantial labeling in the presence of CMP-sia-biotin by western blotting with streptavidin (Suppl. Fig. 4C), and LC-MS/MS proteomic analysis confirmed enrichment of multiple known *O*-glycosylated cell surface/ECM proteins (e.g., CD44, LDLR, FN1, DAG1) including several HSPGs (SDC1, SDC2, SDC4, GPC1, GPC6) (Suppl. Fig. 4D), most of which have been previously identified as *O*-GalNAc modified in other cells and/or tissues in recent studies^23, 40, 41^.

Next, we probed whether targeting *COSMC* or *C1GALT1* alters the secreted proteome in TC28a2 chondrocytes. We collected conditioned media from cultured wild-type, *COSMC^-/-^*, and *C1GALT1^-/-^* cells, respectively, and profiled the secreted proteome by LC-MS/MS by adapting published methods (Fig. 4A) ^42, 43^. This method revealed that truncation of *O*-GalNAc glycans led to remodeling of the secreted proteome, with a total of 62 proteins significantly altered in abundance in *COSMC^-/-^* cells and 92 proteins in *C1GALT1^-/-^* cells, respectively. Notably, we identified 48 differentially secreted proteins shared between both knockout lines (Fig. 4B). Consistent with previous studies^44, 45^, we observed increased secretion of the *O*-glycoprotein LDL receptor (LDLR) in both knockout lines (Fig. 4C-D), with no significant alterations in *LDLR* mRNA expression. The top protein depleted in both knockout lines, the matrix metalloprotease MMP1, exhibited a concurrent decrease in mRNA expression, while additional factors associated with ECM remodeling (MMP9) and chemotaxis (CXCL1, CXCL8) were significantly decreased in the secretome from *COSMC^-/-^*and *C1GALT1^-/-^* cells despite a significant increase in mRNA levels. Several ECM proteoglycans (BGN, HSPG2, COL18A1) showed a reduction at both the transcriptome and secretome levels, while others (AGRN, SRGN) exhibited increased abundance in conditioned media with no corresponding mRNA expression changes. We detected only a single cell surface HSPG (SDC4) in the *C1GALT1^-/-^*secretome, suggesting that shedding of cell surface proteoglycans was minimally impacted in the *COSMC/C1GALT1* knockout cells.

**Figure 4.**
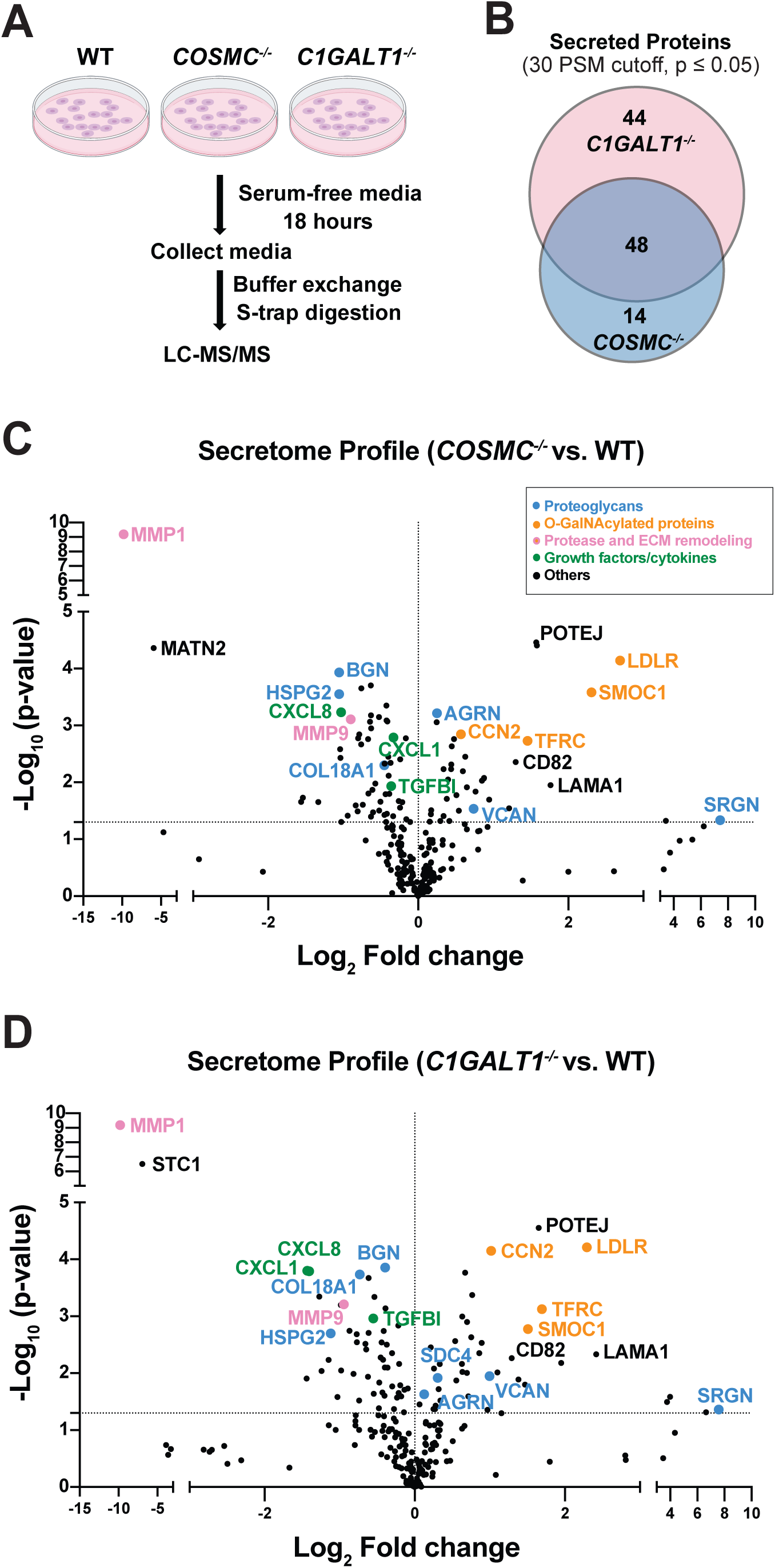
Truncation of *O*-GalNAc glycans leads to remodeling of the secreted glycoproteome in TC28a2 chondrocytes. (A) Workflow for analysis of the secretome from TC28a2 wild-type, *COSMC^-/-^*, and *C1GALT1^-/-^* cells. (B) Venn diagram showing the overlapping protein hits identified in the secretome analysis of *COSMC^-/-^* and *C1GALT1^-/-^* cells versus wild-type controls (n = 3 independent experiments). Over half of the identified protein hits were shared between the knockout cell lines. Volcano plots showing proteins significantly altered in abundance in (C) *COSMC^-/-^*and (D) *C1GALT1^-/-^* cells versus wild-type controls. Proteins were categorized and highlighted as *Proteoglycans*, *O-GalNAcylated proteins*, *Proteases*, *ECM remodeling enzymes*, *Growth factors/cytokines/signaling proteins*, and *other proteins*. p-values were determined by Student t-test, and significantly altered proteins were identified using a threshold of p ≤ 0.05.

### Truncation of *O*-GalNAc glycans reduces cell surface SDC1 via enhanced lysosomal degradation

Among the proteoglycans examined, SDC1 displayed an unexpected discordance between transcript and cell surface abundance. While RNA-seq and secretome proteomics showed no major changes in SDC1 mRNA or shedding, flow cytometry analyses revealed a pronounced decrease in SDC1 cell surface levels (Fig. 3E-F). This discrepancy suggested that truncation of complex *O*-GalNAc glycans might influence SDC1 stability and/or turnover rather than its transcription or shedding. To investigate whether SDC1 trafficking to the cell surface was impaired in COSMC/C1GALT1 mutants, we lifted wild-type or knockout cells with trypsin, respectively, re-plated them, then monitored recovery of SDC1 at the cell surface over time via flow cytometry. SDC1 protein levels were restored to the respective undigested cells after 24 hours at comparable initial rates in both wild-type and knockout cells, despite decreased steady-state SDC1 levels in the COSMC/C1GALT1 deficient lines, indicating that SDC1 trafficking was largely preserved (Fig. 5A-B). Given that SDC1 delivery to the cell surface appeared intact, we next tested whether diminished SDC1 levels in COSMC/C1GALT1 knockout cells resulted from decreased stability and/or increased degradation consequent to the loss of complex *O*-GalNAc glycans. Since SDC1 degradation primarily occurs through endocytosis and lysosomal degradation^46^, we tested whether inhibition of lysosomal activity would restore SDC1 levels in COSMC/C1GALT1 knockout cells. Cells were treated with bafilomycin A1, a V-ATPase inhibitor that impedes lysosomal degradation^47^, and cell surface and total SDC1 levels were monitored via flow cytometry. After a 16-hour incubation with bafilomycin A1, we observed a significant increase in both total and cell surface SDC1 levels in *COSMC^-/-^* and *C1GALT1^-/-^* cells versus vehicle-treated cells (Fig. 5C-E). Confocal imaging corroborated these findings, confirming a complete rescue of SDC1 levels in *COSMC^-/-^* and *C1GALT1^-/-^*cells treated with bafilomycin (Suppl. Fig. 5A-D). Together, these results indicate that loss of complex O-GalNAc glycans destabilizes SDC1 at the cell surface, promoting its endolysosomal-dependent degradation.

**Figure 5.**
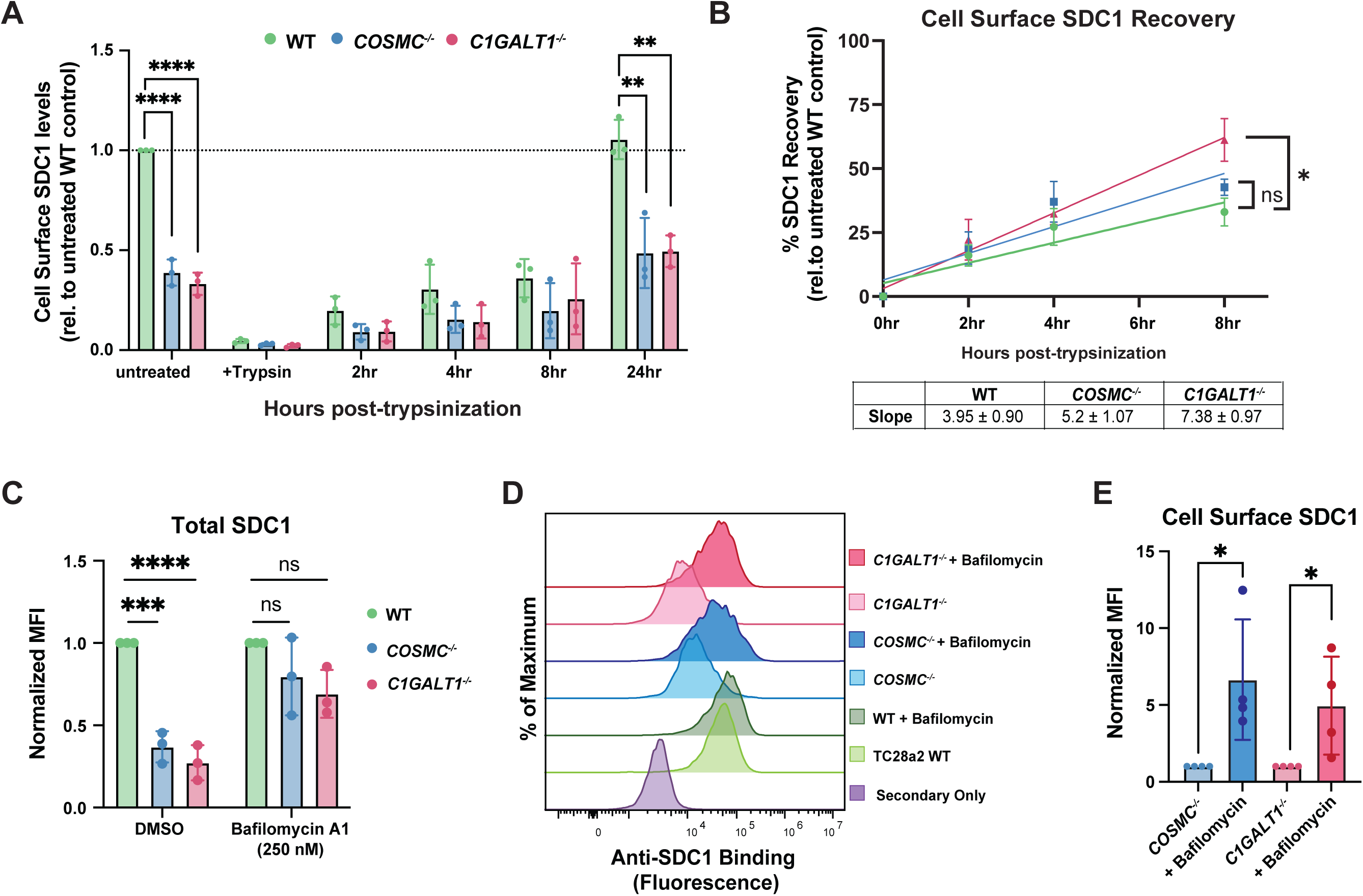
Loss of extended *O*-GalNAc glycans decreases SDC1 cell surface stability and enhances its lysosomal degradation. (A) Time course quantification of cell surface SDC1 levels via flow cytometry prior to and after trypsinization (0-24 hours). Data is normalized to untreated wild-type cells (dashed line). (B) Quantification of SDC1 recovery to the cell surface. Data were normalized to the respective untreated control for each genotype and fitted using linear regression. Table inset includes slopes ± SD for each. Statistical comparisons were performed using an F-test to compare the slopes of knockout lines versus wild-type cells (* p < 0.05). (C) Quantification of total SDC1 protein levels in wild-type, *COSMC^-/-^*, and *C1GALT1^-/-^* cells treated with bafilomycin A1 (0.25 µM) or DMSO vehicle for 16 hours. (D) Representative histograms from flow cytometry analysis of total SDC1 levels in cells treated with bafilomycin A1. (E) Flow cytometry analysis of cell surface SDC1 levels after treatment with bafilomycin A1 or DMSO vehicle. Data points are shown as mean ± SD (n ≥ 3 independent experiments); p-values were determined by one-way ANOVA with Tukey post-test (A,C), and student’s T-test (E) with **** p < 0.0001, *** p < 0.001, ** p < 0.01, * p < 0.05.

### Mucin-type *O*-glycans stabilize cell surface CD44 and regulate CD44v3-FGF1 binding

CD44 is a highly *O*-glycosylated protein^23, 48^ and is a major cell surface receptor on chondrocytes that mediates cell adhesion to the extracellular matrix and participates in matrix organization and turnover^49, 50^. Notably, CD44 can serve as a “part-time” proteoglycan, where a specific CD44 variant, CD44v3, is modified by a single HS chain that can function as a key co-receptor for growth factors, including FGF1^51^ (Fig. 6A). Thus, we investigated whether full length CD44 and/or CD44v3 expression was altered in *COSMC/C1GALT1* knockout cells and whether this may contribute to defects in HSPG presentation and growth factor binding. Despite increased *CD44* transcript levels in both knockout lines (Fig. 3E), immunoblotting and flow cytometry experiments revealed a marked decrease in full-length CD44 protein levels and a lower apparent molecular weight, consistent with loss of complex *O*-glycans (Fig. 6B-C). These results were reproducible in independent *COSMC* and *C1GALT1* knockout clones (Suppl. Fig. 6). Reintroduction of human C1GALT1 into *C1GALT1^-/-^*cells restored cell surface CD44 levels, confirming a direct link between *O*-glycan truncation and CD44 presentation (Fig. 6D). Mechanistically, we also found that cell surface CD44 on COSMC/C1GALT1 deficient cells exhibited increased susceptibility to trypsin-mediated cleavage, with reduced steady-state levels 24 hours post-trypsinization (Suppl. Fig. 6B), indicating that loss of *O*-GalNAc glycans exposes the CD44 ectodomain to enhanced proteolytic turnover at the cell surface. Finally, we also confirmed cell surface expression of CD44v3 in TC28a2 cells using a CD44v3-specific antibody, which showed a marked reduction in both knockout lines (Fig. 6E). To directly assess whether decreased cell surface CD44v3 affects HS-dependent FGF binding, we generated a *CD44^-/-^*knockout line in TC28a2 chondrocytes, validated by Western blot and Sanger sequencing (Fig. 6F, Supplementary Fig. 6C). Both full length CD44 and CD44v3 were depleted in these cells, respectively (Fig. 6F-G). Importantly, *CD44^-/-^* cells exhibited reduced 3G10 and FGF1/FGF2 binding at the cell surface (Fig. 5H-J), closely phenocopying *COSMC^-/-^* and *C1GALT1^-/-^* cell lines and highlighting the important role of CD44 as a cell surface receptor in chondrocytes.

**Figure 6.**
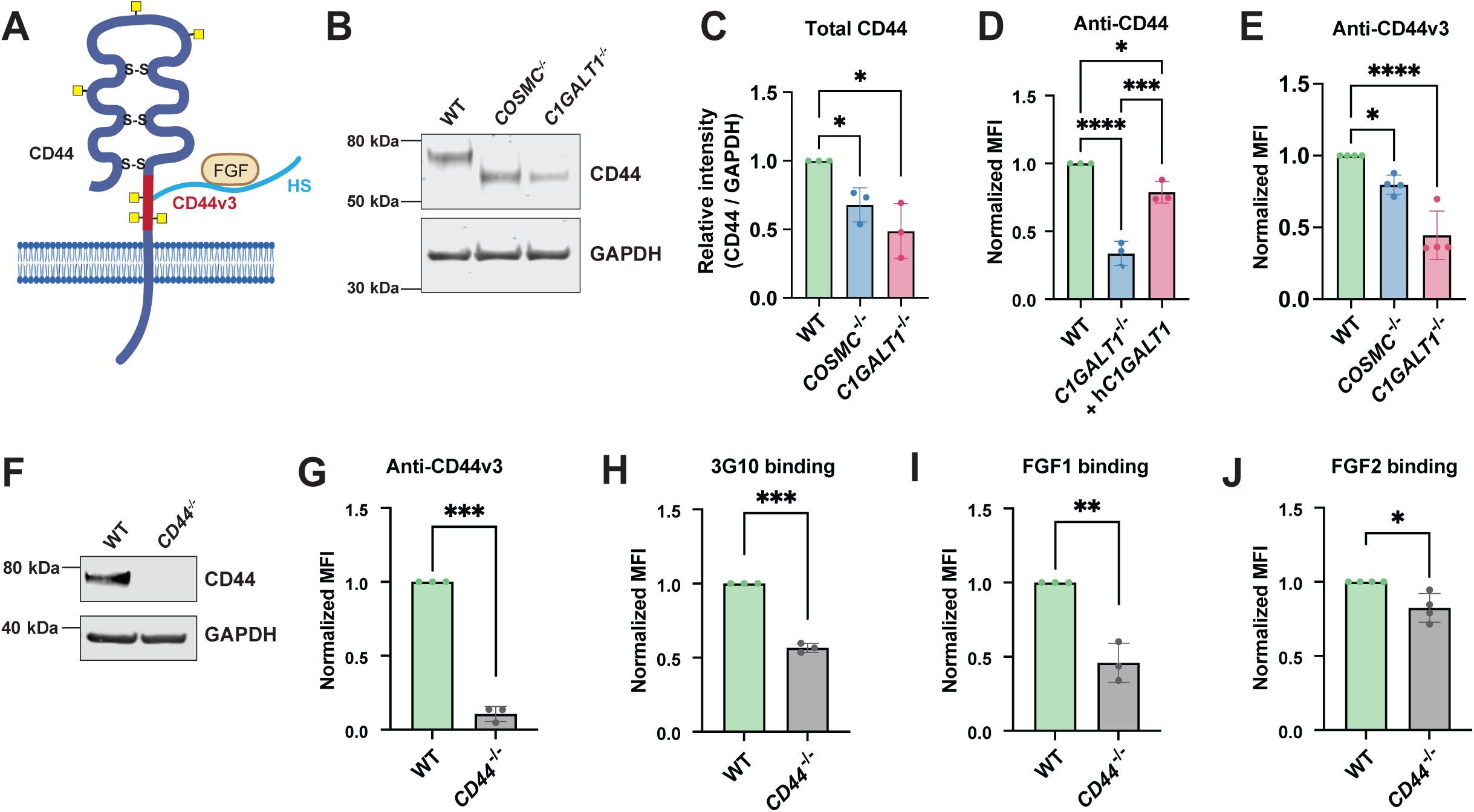
Complex *O*-GalNAc glycans stabilize CD44 and regulate CD44v3-FGF binding. (A) Schematic representation of CD44 protein structure. CD44 is extensively *O*-glycosylated, and alternative splicing generates multiple isoforms, with CD44v3 bearing a heparan sulfate chain that can mediate FGF binding. (B) Western blotting and (C) quantification of CD44 protein levels using a pan-CD44 antibody in TC28a2 wild-type, *COSMC^-/-^*, and *C1GALT1^-/-^* cells, normalized to GAPDH (n = 3 independent experiments). (D) Reintroduction of hC1GALT1 restored cell surface CD44 levels in *C1GALT1^-/-^* knockout cells. (E) Detection and quantification of cell surface CD44v3 levels in wild-type and mutant lines using an anti-CD44v3 antibody and flow cytometry. (F) Validation of TC28a2 *CD44^-/-^* knockout cells via Western blot using a pan-CD44 antibody. Flow cytometry analyses of cell surface CD44v3 levels, (H) 3G10 binding, and (I) FGF1 and (J) FGF2 binding in wild-type and *CD44^-/-^* cells. Data points are shown as mean ± SD (n ≥ 3 independent experiments); p-values were determined by one-way ANOVA with Tukey post-test (C-E), and student’s T-test (G-J) with **** p < 0.0001, *** p < 0.001, ** p < 0.01, * p < 0.05.

### Ablation of *Cosmc/C1galt1* impairs chondrogenic maturation and cartilage matrix formation in growth plate-like chondroprogenitors

Given our findings that truncation of *O*-GalNAc glycans destabilizes cell surface proteoglycans and impairs cell signaling, we next investigated whether mucin-type *O*-glycosylation influences chondrocyte differentiation. To test this, we employed CRISPR/Cas9 to ablate *Cosmc* or *C1galt1* in murine growth plate-like chondroprogenitors (GPLCs), an established in vitro model of the epiphyseal growth plate^52, 53^. Clonal knockout lines were validated by Sanger sequencing and VVA/PNA lectin binding (Fig. 7A-B, Suppl. Fig. 7A). *Cosmc^-/-^* and *C1galt1^-/-^* cells exhibited significantly reduced 3G10 staining and diminished cell surface binding of FGF1 and FGF2 (Fig. 7C-E), accompanied by reduced CD44 (Fig. 7F), mirroring our results in human TC28a2 chondrocytes. When cultured in high-density micromass and stimulated with insulin-transferrin-selenium (ITS) to promote chondrogenic maturation^54^, *Cosmc^-/-^* and *C1galt1^-/-^* cells exhibited significantly reduced Alcian blue staining at Day 7 (Fig. 7G-H), indicating impaired glycosaminoglycan-rich cartilage matrix deposition^55^. Furthermore, expression of key matrix genes, *Acan* and *Col2a1*^56^, was reduced (Fig. 7I-J), while *Sox9* expression remained unchanged (Fig. 7K), suggesting that lineage commitment was preserved but downstream matrix elaboration and maturation were impaired. Overall, these data suggest that truncation of *O*-GalNAc glycans disrupts proteoglycan stability and HS-dependent signaling necessary for chondrogenic maturation.

**Figure 7.**
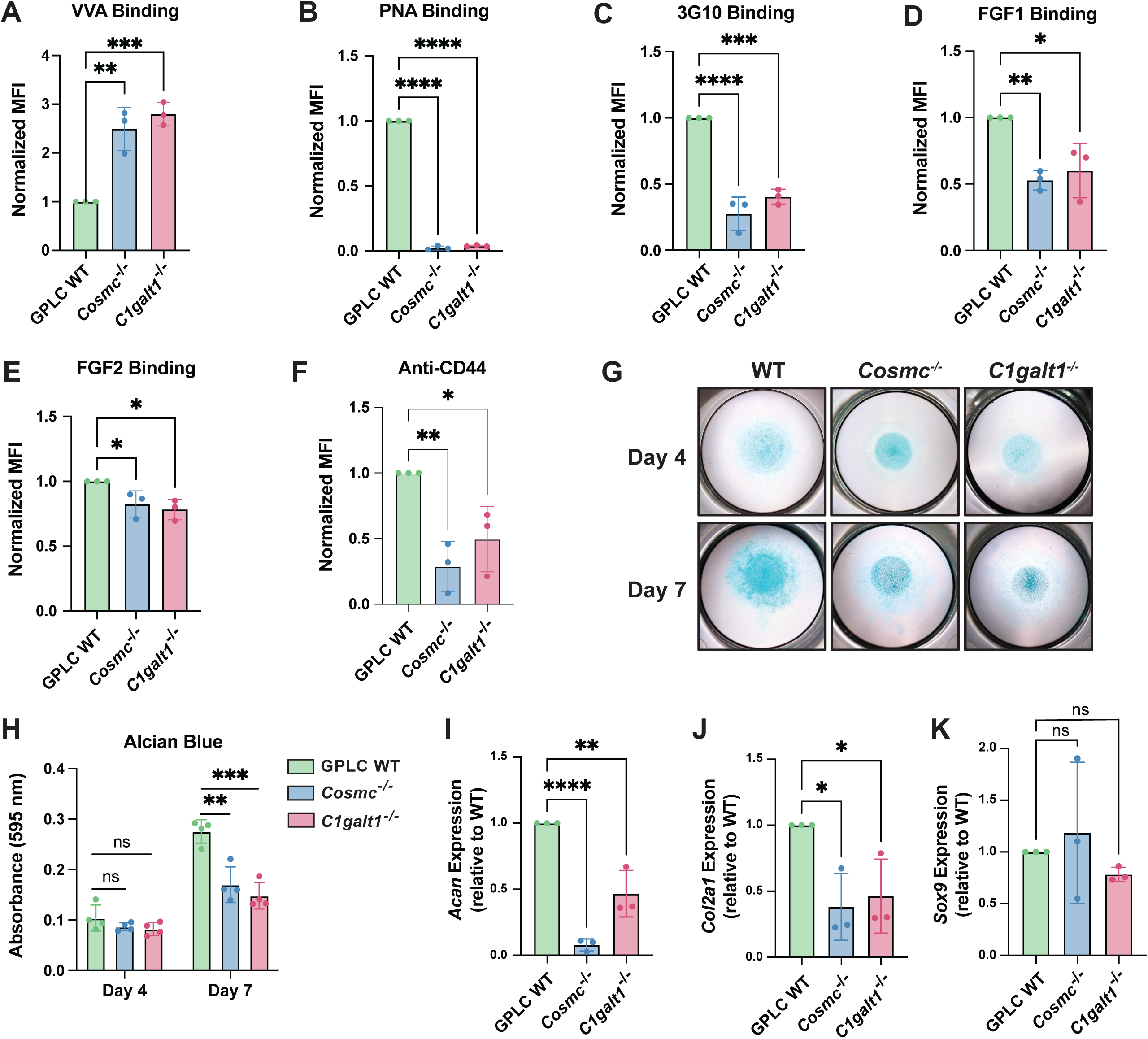
Disruption of *O*-GalNAc glycosylation impairs chondrogenic maturation and cartilage matrix elaboration in growth plate-like chondroprogenitors. Flow cytometry analysis of (A) VVA lectin, (B) PNA lectin, (C) 3G10 antibody, (D) FGF1, (E) FGF2, and (F) pan-CD44 antibody binding in wild-type and knockout growth-plate-like chondroprogenitors (GPLCs). (G) Micromass cultures of GPLC wild-type, *Cosmc^-/-^*, and *C1galt1^-/-^* cells at day 4 and day 7 after chondrogenic induction with insulin-transferrin-selenium (ITS). At each time point, cultures were fixed and stained with Alcian blue. (H) Quantification of Alcian blue staining confirmed visual data and confirmed statistical significance at Day 7. qPCR analysis of chondrogenic markers (I) *Acan* (J) *Col2a1*, and (K) *Sox9* in wild-type and knockout GPLC micromass cultures at Day 7. Data points are shown as mean ± SD (n ≥ 3 independent experiments); p-values were determined by one-way ANOVA with Tukey post-test with **** p < 0.0001, *** p < 0.001, ** p < 0.01.

## Discussion

In this study, we report that complex mucin-type *O*-glycosylation regulates proteoglycan expression, stability, and function. Through CRISPR-mediated genetic ablation of *COSMC*/*C1GALT1*, transcriptomic and proteomic profiling, and functional analyses, we show that loss of extended *O*-GalNAc glycans impairs cell surface presentation of SDC1 and CD44, limits heparan sulfate presentation, and attenuates growth factor-dependent signaling and chondrogenesis. These findings uncover a mechanistic link between *O*-GalNAc glycan maturation and proteoglycan biology, revealing that crosstalk between glycosylation pathways contributes to proper extracellular matrix organization and cartilage differentiation.

Our studies intriguingly identified a selective reduction in cell surface HSPG presentation in chondrocytes following loss of extended core 1 *O*-glycans, with seemingly minimal effects on HS and CS/DS composition (Fig. 1, Suppl. Fig. 1). The specificity of this effect would suggest that *O*-GalNAcylation preferentially influences the biosynthesis or stability of HSPG core proteins rather than global glycosaminoglycan chain synthesis. The ∼6-fold higher prevalence of HS chains versus CS/DS at the cell surface of TC28a2 chondrocytes (Fig. 1F-G) likely drives the observed impact of truncated *O*-GalNAc glycans on HSPG presentation. There are likely differential effects on proteoglycan presentation in different cell types and tissues, largely dependent on gene expression profile and site-specific glycosylation of proteoglycans. Notably, our work also highlighted differential effects on proteoglycan secretion in *COSMC/C1GALT1* knockout cells (Fig. 4), with altered abundance of certain ECM HSPGs (HSPG2, COL18A1, AGRN, SRGN) and CSPGs (BGN, VCAN), suggesting that extended *O*-GalNAc glycans likely regulate both proteoglycan secretion and their cell surface stability. The observed impact on shedding and secretion of ECM proteins is supported by previous studies implicating mucin-type *O*-glycans as regulators of protein trafficking and secretion^57, 58, 59, 60^. Our results extend this paradigm by showing that *O*-GalNAc truncation impairs the stability and persistence of HSPGs, including SDC1 and the “part-time” proteoglycan CD44v3, thereby altering the abundance and organization of HS chains at the cell surface in chondrocytes.

A previous study reported that chondrocyte-specific overexpression of *Galnt3*, a ppGalNAc-transferase, increased Tn-antigen levels and reduced GAG production in mice and resulted in defects in chondrocyte organization, hypertrophy, and skeletal development^18^, suggesting that *O*-GalNAc initiation influences proteoglycan synthesis and function. It is possible that overexpression and/or knockout of *Galnt3* leads to competition of *O*-glycan initiation at serine residues within the proteoglycan core, where xylosyltransferases (XYLT1/XYLT2) can add xylose to initiate GAG assembly (Fig. 1H). The site-specific regulation of diverse *O*-glycan modifications on their respective protein attachment sites is still largely unknown, with evidence of amino acid sequence^61^ and/or substrate specificity of the enzymes^24^ likely contributing to *O*-glycan initiation. Enrichment of VVA lectin staining and Tn antigen presentation reported in *Galnt3* transgenic mice^18^ mimics the observed phenotypes in *COSMC/C1GALT1* deficient chondrocytes generated in the current study. Together, these findings suggest that truncation of *O*-GalNAc glycans contributes to reduced proteoglycan expression and destabilization of cell surface PGs, potentially leading to skeletal defects at the growth plate. Future studies will investigate how conditional ablation of C1GALT1/COSMC at the growth plate in animal models impacts skeletal development. The results presented in the current study reveal distinct mechanisms by which loss of complex *O*-GalNAc glycans alters proteoglycan dynamics, including transcriptional downregulation of genes encoding proteoglycans (Fig. 3) and destabilization of HSPGs at the cell surface. Specifically, despite upregulation of *SDC1* and *CD44* mRNA expression in *COSMC/C1GALT1* knockout cells, we found that both these proteins, which are both highly *O*-GalNAcylated^23, 37^, required complex mucin-type *O*-glycosylation for proper cell surface presentation on chondrocytes. Reduced SDC1 protein levels were caused by enhanced lysosomal degradation (Fig. 5) rather than decreased synthesis or enhanced shedding/secretion (Figs. 3-4), and decreased CD44 levels were likely caused by increased susceptibility to proteolysis upon loss of extended *O*-GalNAc glycans (Suppl. Fig. 6), consistent with prior studies^48^. In addition, loss of a specific HS-modified CD44 variant, CD44v3, contributed to reduced HS-protein interactions at the cell surface (Fig. 6). Overall, these findings shed light on the multi-functional roles of mucin-type *O*-glycans in regulation of proteoglycan stability and function.

Clinically, CDG disorders often present with skeletal abnormalities, including *COSMC-CDG*^15^ and MHE^17^. Our in vitro chondrogenesis model of the growth plate revealed that truncation of *O*-GalNAcylation led to decreased chondrogenic differentiation and reduced expression of key cartilage matrix genes (Fig. 7). Since HSPGs are crucial for bone development, genetic defects in mucin-type *O*-glycan assembly would negatively impact HSPG presentation and function, thus contributing to skeletal and cartilage defects found across CDG disorders. Interestingly, *COSMC-CDG* patient-derived fibroblasts, which harbor a hypomorphic mutation in *COSMC* and exhibit reduced T-synthase activity^15^, also showed reduced cell surface HSPGs and FGF binding (Fig. 2I), suggesting other cell types may also be similarly impacted. Therapeutically, restoring HSPG function or supplementation of functional HSPGs may represent promising strategies to mitigate selected clinical manifestations associated with impaired glycosylation in patients. Beyond CDGs, truncated *O*-GalNAc glycans are a hallmark found in multiple cancers, where the Tn antigen regulates tumor cell proliferation, differentiation, and other oncogenic features^8, 62, 63^. Cancer progression also depends on proteoglycan function^64^; thus, regulation of mucin-type *O*-glycosylation may also alter proteoglycan abundance and composition in tumor cells, potentially aiding tumor invasion and metastasis.

Collectively, our results support a model in which extended *O*-GalNAc glycans stabilize proteoglycan core proteins, preserve HS chain presentation, and maintain FGF-FGFR signaling essential for cartilage development (Fig. 8). More broadly, our findings suggest that glycan cross-regulation plays a crucial role in cellular function, development, and disease progression. These findings underscore the interplay between *O*-GalNAcylation and proteoglycan structure and function and illustrate the role of diverse glycosylation pathways in shaping the cell surface proteome. Future studies will define how specific *O*-GalNAc sites on proteoglycan core proteins regulate their folding, export, and stability, and whether targeted manipulation of *O*-glycan processing can restore proteoglycan function in disease contexts. Given the emerging evidence of glycan interdependence across cell types and tissues, elucidating these mechanisms will deepen our understanding of glycosylation crosstalk and may identify new therapeutic opportunities for glycosylation disorders.

**Figure 8.**
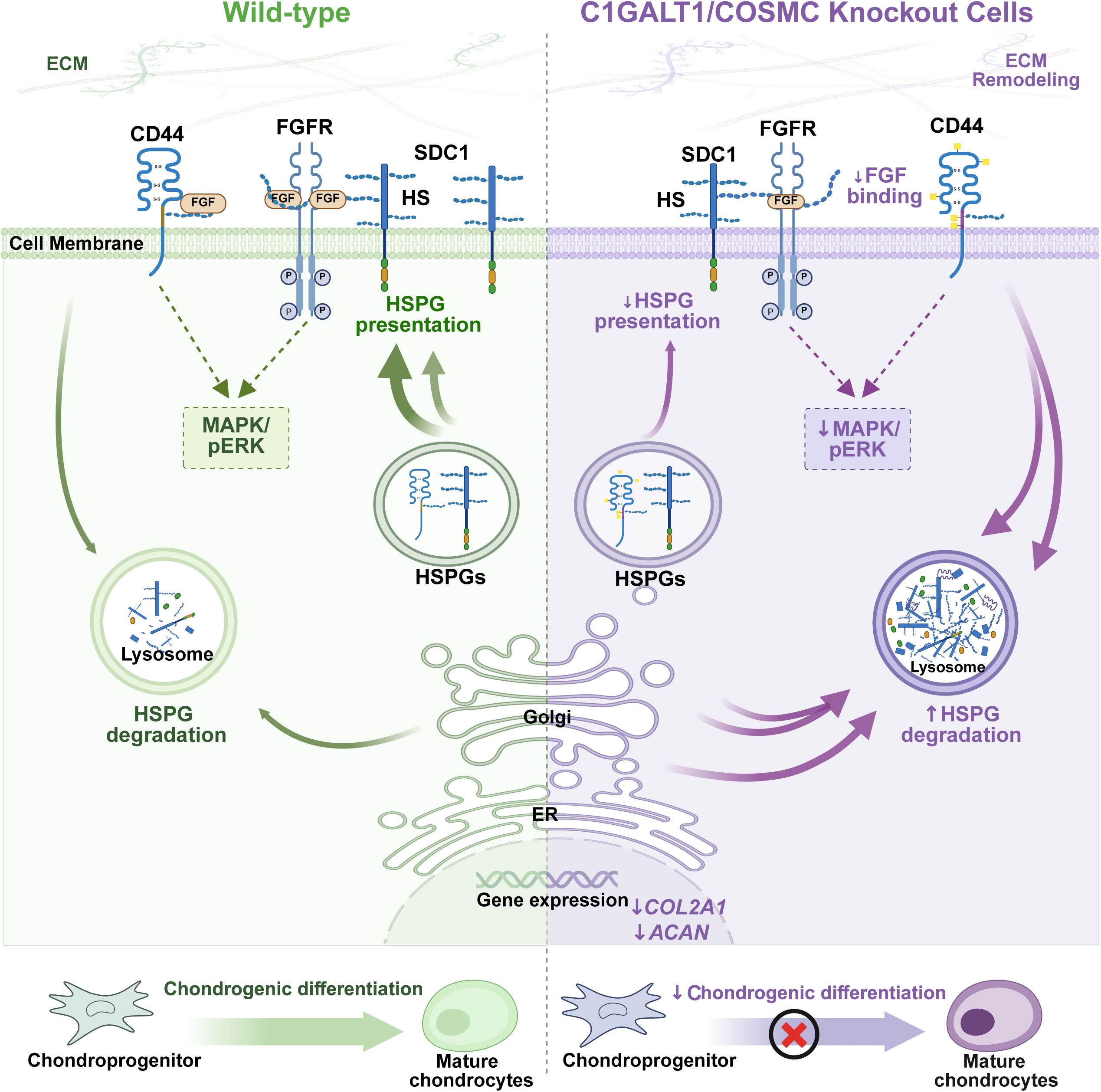
Proposed model of proteoglycan homeostasis and ECM remodeling alterations caused by truncation of *O*-GalNAc glycans in *COSMC/C1GALT1*-deficient cells.

## Materials and methods

### Cell culture

TC28a2 (Sigma-Aldrich #SCC042) and HEK293T cells (ATCC CRL-3216) were cultured in DMEM (Gibco) with 10% (v/v) FBS (Gibco) and 1% (v/v) penicillin/streptomycin at 37 °C under an atmosphere of 5% CO2/95% air. The growth plate like chondrocyte (GPLC) cell line, derived and v-myc immortalized from 13DPC mouse limb buds as published^52^, was obtained as a gift from Dr. Vicki Rosen (Harvard School of Dental Medicine) and were grown in DMEM/F12 (Gibco) with 10% (v/v) FBS (Gibco) and 1% (v/v) penicillin/streptomycin at 37°C with 5% CO2/95% air. Patient-derived fibroblasts from a male *COSMC-CDG* patent (A20D-COSMC, M2)^15^ were obtained as a gift from Dr. Richard Cummings (Harvard Medical School) and were grown in DMEM (Gibco) with 10% (v/v) FBS and 1% (v/v) penicillin/streptomycin at 37°C with 5% CO2/95% air. For routine passaging, all cells listed above were detached using 0.05% trypsin-EDTA (Thermo), sub-cultured every 3 to 4 days, and revived from liquid nitrogen after ≤ 10 passages.

### Cell line generation

HEK293T cells were co-transfected with Fugene 6 (Promega) and 4 μg each of a viral envelope plasmid (pMD2.g, a gift from Didier Trono and purchased from Addgene, #12259), packaging plasmid (psPAX2, a gift from Didier Trono and purchased from Addgene, #12260), and Cas9 expression plasmid (lentiCas9-Blast, a gift from Feng Zhang and purchased from Addgene, #52962) to generate Cas9 lentiviral particles. Lentiviral particles were collected from HEK293T cells and filtered through a 0.45 μm syringe filter to remove cell debris before use in subsequent transductions. Cas9 lentiviral particles were added to TC28a2 or GPLC wild-type cells, followed by selection with blasticidin (2 μg/mL) to generate Cas9-expressing cells. To generate *COSMC*, *C1GALT1*, and *CD44* knockout cell lines, sgRNAs sequences were cloned into the lentiGuide-puro vector (a gift from Feng Zhang and purchased from Addgene, #52963). These constructs were co-transfected with Fugene 6 into HEK293T cells along with viral plasmids pMD2.g and psPAX2 to produce lentiviral particles, which were subsequently used to transduce Cas9-expressing TC28a2 and GPLC cells, respectively. All sgRNA targeting sequences, and primers used for genotyping, are listed in Supplementary Tables 1-2. After selection of the transduced cell pool with puromycin (0.2 μg/mL) for at least 4 days, surviving cells were seeded onto a 96-well plate by limiting dilution to establish clonal populations.

### Western blotting

Total protein was extracted from cells using RIPA lysis buffer (EMD Millipore) supplemented with protease inhibitors (Roche). Protein concentrations were determined by Bicinchoninic Acid (BCA) assay (Thermo Scientific-Pierce). Equal amounts of protein were loaded and separated on SDS-PAGE gels (4-12% Bis-Tris, NuPAGE, Invitrogen), transferred onto nitrocellulose membranes (Thermo), and blocked with Intercept Blocking Buffer (LI-COR Biosciences) for 1 hour at room temperature. Membranes were probed with primary antibodies against COSMC (Proteintech, #19254-1-AP; 1:1,000), C1GALT1 (Santa Cruz Biotechnology, #sc-100745; 1:500), CD44 (Cell Signaling Technology, #3570; 1:1000), p44/42 MAPK (ERK1/2; Cell Signaling Technology, #9107; 1:1000), phospho-p44/42 MAPK (p-ERK; Cell Signaling Technology, #4370, 1:2000), GAPDH (Cell Signaling Technology, #5174; 1:2500), or β-actin (Cell Signaling Technology, #3700; 1:5000) at 4°C overnight. After washing with TBST, mouse and rabbit primary antibodies were incubated with secondary Odyssey IR dye antibodies (anti-mouse 800, LI-COR Biosciences, #926-32212; donkey anti-rabbit 680, LI-COR Biosciences, #926-68073; 1:14,000) and visualized on an Odyssey infrared imaging system (LI-COR Biosciences). Band intensities were quantified using ImageJ and normalized to housekeeping proteins.

### MAPK/ERK signaling assays

For signaling assays, wild-type or knockout TC28a2 cells were cultured in 12-well plates. The growth medium was exchanged for serum-free DMEM (Invitrogen) for 5 hours followed by the addition of FGF2 (bFGF, Peprotech, 0- 10 ng/mL) for 5 min before placing the cells on ice. Next, the cells were washed twice with ice-cold PBS and were lysed using RIPA buffer containing protease inhibitors. Protein concentration was determined using the Pierce BCA Protein Assay Kit (Thermo Scientific) before Western blot analysis. Cells were resolved by SDS-PAGE and blotted with primary antibodies against p-ERK and ERK antibodies (Cell Signaling Technology. Bands were visualized on an Odyssey Infrared imaging system (Li-Cor Biosciences). Band intensities were quantitated by densitometry using Image J software.

### Protein biotinylation

Heparin-Sepharose (100 μl, Cytiva) was pre-equilibrated with PBS (Gibco) and then loaded with human FGF1 (Peprotech, #100-17A) or human FGF2 (Peprotech, #100-18B) dissolved in PBS, as previously described^65^. The flow-through was collected and reloaded onto the column twice to ensure complete binding. After washing twice with PBS, 0.6 mg/mL of Sulfo-NHS-LC-biotin (Thermo Fisher) in PBS was applied to the column and incubated for 1 hour at room temperature. Each column was washed three times with PBS, then bound biotinylated protein was eluted with 0.4 mL of PBS buffer containing an additional 2 M NaCl. All biotinylated proteins were stored at - 80 °C before use.

### Flow cytometry and cell surface binding experiments

TC28a2 and GPLC cells were grown in monolayer culture, washed twice with PBS, and detached using Accutase (Sigma-Aldrich, #A6964) via incubation for 5 min at room temperature. Cells were then resuspended in PBS + 0.1% BSA (FACS buffer). For Tn and T antigen detection, 1 × 10^6^ cells were pre-incubated with sialidase (Lectenz Bio, #GE0701; 50 U per 100 µL) for 1 h at 37 °C, washed with FACS buffer, and stained with the VVA-FITC (Vector Labs, #FL-1231-2, 40 ng/mL), anti-Tn antibody (Thermo Scientific, #MA1-80055, 1 μg/mL), or PNA lectin tagged with Alexa Fluor 647 (Thermo, #L32460, 20 ng/mL) for 45 min on ice in the dark.

For the detection of other proteins and ligands, lifted cells were incubated in suspension for 30 min at 4 °C with 0.5 μg/mL mAb 10E4 (AMSBio #370255-1, Clone F58-10E4, 1:5000), 1 μg/mL mAb 3G10 (AMSBio, #370260-S, clone F69-3G10, 1:1000), 1 μg/mL mAb 2B6 (AMSBio # 270432-CS, Clone 2B6, 1:200), 0.1 μg/mL pan-CD44 mAb (Thermo #14-0441-82, Clone IM7, 1:1000), 0.5 μg/mL anti-CD44v3 mAb (R&D Systems #BBA11, Clone 3G5, 1:1000), 0.2 μg/mL anti-SDC1 mAb (Novus Biologicals #NB100-64980, Clone B-A38, 1:500), 0.5 μg/mL anti-SDC2 mAb (R&D Systems #MAB2965, Clone 305515, 1:1000), 80 nM biotin-FGF1, or 2.5 nM biotin-FGF2, respectively. Where applicable, cells were pre-treated with 5 mU/mL each of heparin lyases I, II, and III (BBI Solutions) for 30 min at 37 °C to generate HS neoepitopes for 3G10 binding, or with 2 mU/mL chondroitinase ABC (Sigma-Aldrich, #C3667-5UN) for CS neoepitope detection for 2B6 binding. Bound 10E4 was detected with 2 μg/mL anti-mouse IgM AlexaFluor 647 (Invitrogen, #A-21238, 1:1000). Primary IgG antibodies were subsequently detected with 2 μg/mL anti-mouse IgG AlexaFluor 647 (Invitrogen, #A21235, 1:1000), 2 μg/mL anti-mouse IgG AlexaFluor 488 (Invitrogen, #A11001, 1:1000) or 2 μg/mL anti-rat IgG AlexaFluor 488 (Invitrogen, #A11006, 1:1000). Binding of biotinylated proteins was detected by streptavidin-Cy5 (Molecular Probes, 1:1000). Flow cytometry was performed using a CytoFLEX S instrument (Beckman Coulter; ≥10,000 events/sample). Live cells were gated based on forward and side scattering. The extent of protein binding was quantified using the geometric mean of the fluorescence intensity. Unstained negative controls were included in each experiment. Data were plotted and analyzed in GraphPad Prism v10.8.

For quantification of total SDC1 protein levels by flow cytometry, monolayer cultures were detached with Accutase (Sigma-Aldrich, A6964; 1 mL per 10-cm dish), fixed using 4% paraformaldehyde for 20 min at room temperature, washed twice with PBS, and permeabilized with 1% Triton X-100 for 5 min. After a DPBS wash, cells were blocked with 2% BSA in PBST on ice for 30 min and then incubated with primary antibody against 0.2 µg/mL SDC1 (Fisher Scientific, N106491010U; 1:500). After washing, cells were labeled with 2µg/mL Alexa Fluor 647-conjugated anti-mouse IgG secondary antibody (Fisher, A11001; 1:1000) and analyzed via flow cytometry following standard procedures.

### Proteoglycan cell surface recovery assays

To assess recovery of cell surface SDC1 and CD44, TC28a2 wild-type, *COSMC*^⁻/⁻^, and *C1GALT1*^⁻/⁻^ chondrocytes were plated and grown to confluence. Cells were then detached by incubation with 0.25% trypsin-EDTA (Thermo, #25200056) for 15 min at 37 °C. After neutralization and resuspension, 2 × 10^6^ cells were replated and allowed to recover for the indicated times (0, 2, 4, 8, and 24 hours post-trypsinization). Untreated wild-type and knockout cells were maintained in parallel and served as non-trypsinized controls. At each time point, cells were lifted with Accutase, fixed in 4% paraformaldehyde, and analyzed for cell surface SDC1 or CD44 via flow cytometry using the antibodies and methods described above. For bafilomycin A1 experiments, cells were pre-treated with bafilomycin A1 (0.25 µM) or vehicle (DMSO) for 16 hours prior to flow cytometry or immunofluorescence analyses.

### GAG isolation, purification and LC-MS disaccharide analysis

2×10^6^ cells were seeded into a 15 cm plate and harvested when confluent. Briefly, cells were washed with PBS, lifted with trypsin (Gibco), and the trypsin-released glycosaminoglycans were digested with Pronase (0.5 mg/mL, Sigma) overnight at 37 °C. The product was filtered and passed through a DEAE-Sephacel (Cytiva) column equilibrated in 50 mM sodium acetate buffer, pH 6.0, containing 200 mM NaCl then passed through a PD-10 desalting column (Cytiva). For HS disaccharide analysis, lyophilized GAGs were incubated with 2 mU each of heparin lyases I, II, and III for 16 h at 37 °C in a buffer with 40 mM ammonium acetate and 3.3 mM calcium acetate, pH 7. For CS analysis, lyophilized GAGs were incubated with 50 mM Tris and 50 mM NaCl, pH 7.9. HS and CS disaccharides were aniline-tagged and analyzed via GRIL-LC/MS, using established methods^26, 66^. All LC-MS data analysis and results are included in Supplementary Tables 4-9 in the supplementary information

### C1GALT1 rescue experiments

The sgRNA-resistant C1GALT1 cDNA vector was generated by introducing silent mutations into the sgRNA target site (5’-TTTAAGCCTTATGTAAAGCA-3’ mutated to 5’-TTTAAGCCTTATGTTAAACA-3’) in a C1GALT1 expression vector (Sino Biological, # HG19234-UT). C1GALT1 cDNA was then subcloned into a pLX208-CMV-Hygro plasmid (Addgene #153007). Lentiviral particles were produced in HEK293T cells by co-transfection with the expression plasmid, psPAX2 packaging plasmid (Addgene #12260), and the VSV-G encoding plasmid pMD2.G (Addgene #12259), as described above. Viral supernatant was collected and used to transduce TC28a2 *C1GALT1*^-/-^ knockout cells. Stably infected cells were selected with hygromycin (75 µg/mL) for 5 days, and C1GALT1 expression was confirmed by immunoblotting.

### RNA extraction, qPCR, and RNA sequencing

Total RNA was isolated from cells using TRIzol (Invitrogen, 15596026) and the RNeasy Kit (Qiagen), following the manufacturers’ instructions. cDNA synthesis was carried out with the SuperScript IV First Strand Synthesis Kit (Invitrogen) using random hexamers, according to the manufacturer’s guidelines. Quantitative PCR was performed with cDNA and SYBR Green Master Mix (Applied Biosystems) following the manufacturer’s protocols. The expression levels of *YWHAZ* or *Gapdh* were used to normalize the target gene expression across human and mouse samples, respectively. The primers used for gene expression analyses are listed in Supplementary Table 3.

### RNA sequencing and differential gene expression analysis

For transcriptome analysis, total RNA from wild-type and knockout cell lines was used for library preparation and next-generation sequencing (IGM Genomics Core, UC San Diego). Raw sequencing reads were processed with the GeneGlobe RNA-seq Analysis Portal (Qiagen) to remove adapters and low-quality bases, then aligned to the human genome (GRCh38; GCF_000001405.38). Differential gene expression analysis was conducted within the same portal, considering genes with an adjusted p-value ≤ 0.01 and a fold change of ± 2 as differentially expressed. Functional annotation and gene set enrichment analysis of the top differentially expressed genes were performed using Metascape (http://metascape.org/)^67^.

### Expression and purification of human ST6GALNAC1

The expression construct for human ST6GALNAC1 was generated encoding the truncated domain of human ST6GALNAC1 (UniProt Q9NSC7, residues 36-600) as an NH2-terminal fusion protein in the pGEn2 expression vector, as described in prior studies^68^. Briefly, the fusion protein coding region was comprised of a 25-amino acid signal sequence, an His8 tag, AviTag, the “superfolder” GFP, the 7-amino acid recognition sequence of the tobacco etch virus (TEV) protease followed by the truncated ST6GALNAC1 coding region. The ST6GALNAC1 expression construct was used for transient transfection of suspension culture HEK293-F cells (FreeStyle^TM^ 293-Fcells, Thermo Fisher Scientific) maintained at 0.5-3.0x10^6^ cells/mL in a humidified CO2 platform shaker incubator at 37°C with 50% relative humidity and 125 RPM. Transient transfection was performed using HEK293-F cells in the expression medium comprised of a 9:1 ratio of Freestyle^TM^293 expression medium (Thermo Fisher Scientific) and EX-Cell expression medium including Glutmax (Sigma-Aldrich). Transfection was initiated by the addition of plasmid DNA and polyethyleneimine as transfection reagent (linear 25-kDa polyethyleneimine, Polysciences, Inc.). Twenty-four hours post-transfection the cell cultures were diluted with an equal volume of fresh media supplemented with valproic acid (2.2 mM final concentration) and protein production was continued for an additional 5 days at 37°C, 125 RPM and 5% CO2. Cell cultures were harvested, clarified by sequential centrifugation at 1200 RPM for 10 min and 3500 RPM for 15 min at 4°C, and passed through a 0.8 µM filter (Millipore, Billerica, MA). The protein preparations were adjusted to contain 20 mM HEPES, 20 mM imidazole, 300 mM NaCl, pH 7.5, and subjected to Ni-NTA Superflow (Qiagen, Valencia, CA) chromatography using a column preequilibrated with 20 mM HEPES, 300 mM NaCl, 20 mM imidazole, pH 7.5 (Buffer I). Following loading of the sample, the column was washed with 3 column volumes of Buffer I followed by 3 column volumes of Buffer I containing 50 mM imidazole and eluted with Buffer I containing 300 mM imidazole at pH 7.0. The protein was concentrated to ∼3 mg/mL using an ultrafiltration pressure cell (Millipore) with a 10-kDa molecular mass cutoff membrane and further purified by size exclusion chromatography using superdex 200 column, peak fractions were pooled and buffer exchanged with 20 mM HEPES, 150 mM NaCl, pH7.0, 10% glycerol and adjusted at 1 mg/mL concentration. The final protein preparation was aliquoted and stored at -80°C until use.

### Expression and purification of Sialidase from *Paenarthrobacter ureafaciens*

The expression cassette for sialidase from *Paenarthrobacter ureafaciens* (Uniprot Q5W7Q2: residues 39-424) was generated in the pET vector. Briefly the cassette was comprised of a “superfolder” GFP, an His6 tag, followed by Sialidase coding region. The protein expression was carried out in arctic express using IPTG induction (0.1 mM) overnight at 18°C. The cell pellets were harvested at 8000 x g for 20 min at 4°C and further subjected for purification. The cell pellet (∼5 gm) was resuspended in 20-25 mL of binding buffer I (20 mM HEPES, 300 mM NaCl, 20 mM imidazole, 5% glycerol, pH 7.5). The lysis was carried out using an ultrasonication probe (30 second pulse at an amplitude of 50 followed by cooling on ice for 1min) and the sonication procedure was repeated two times (total sonication time was 6 min in each cycle). After sonication, cell debris was removed by centrifugation (4°C, 12000 rpm, 30 min). The clear supernatants were then diluted ten times in binding buffer and loaded on Ni-NTA superflow (Qiagen, Valencia, CA) chromatography column. Following loading of the sample, the column was washed with 3 column volumes of Buffer I followed by 3 column volumes of Buffer I containing 50 mM imidazole, and eluted with Buffer I containing 300 mM imidazole at pH 7.0. The protein was concentrated to ∼3 mg/mL using an ultrafiltration pressure cell (Millipore, Billerica, MA) with a 10-kDa molecular mass cutoff membrane and further purified by size exclusion chromatography using superdex 75 column, peak fractions were pooled and buffer exchanged with 20 mM HEPES, 100 mM NaCl, pH7.0, 10% glycerol, 0.05% sodium azide and adjusted at 1 mg/mL concentration. The final protein preparation was aliquoted and stored at -80°C until use.

### Neuraminidase-coupled selective exoenzymatic labeling with ST6GALNAC1

SEEL was utilized as previously described^38^, with modifications. TC28a2 wild-type, *COSMC*^-/-^, and *C1GALT1*^-/-^ cells were cultured to confluence in 10-cm dishes and washed three times with 5 mL PBS to remove residual serum-containing media. Cells were detached by adding PBS and gently scraping with a cell scraper. Cells were pelleted and incubated for 2 hours at 37°C with end-over-end rotation in 600 µL of serum-free DMEM containing recombinant ST6GALNAC1 (42 µg/mL), CMP-Neu5Ac-C5-triazole-biotin (100 µM, synthesized as reported^38^), BSA (2 mg/mL, 2 µL per 300 µL reaction), alkaline phosphatase (FastAP, Thermo Scientific, #EF0651; 2 µL per 300 µL reaction), and sialidase (10 µL per 300 µL). Negative control samples lacked CMP-Neu5Ac-C5-triazole-biotin treatment. Cells were washed three times with 1 mL of PBS and lysed in RIPA buffer containing protease inhibitors (Thermo Scientific, #88666). Cell lysates were clarified by centrifugation, protein concentration was measured by the BCA assay, incubated overnight at 4°C with end-over-end rotation, and adjusted for amount of Streptavidin agarose beads (Thermo Scientific) to enrich biotinylated proteins. Streptavidin beads were collected on a magnetic stand, supernatants were retained as flow-through, and beads were washed extensively including two 1 mL washes with RIPA buffer, 1 mL high salt wash with 1M KCl in 25 mM Tris (pH 7.4), 1 mL high pH wash with 0.1 M Na2CO3 in MilliQ water, 1 mL with 2 M urea in TBS (pH 8), and three 1 mL washes with RIPA buffer (1X TBS washes were included between each listed wash condition). Bound proteins were eluted in 50 mM triethylammonium bicarbonate (Sigma) containing 5% sodium dodecyl sulfate (Sigma) at 95°C and subjected to protein digestion and processing for LC-MS/MS analysis. Eluted proteins were reduced by incubation with 10 mM of dithiothreitol (Sigma) at 56°C and alkylated by 27.5 mM of iodoacetamide (Sigma) at room temperature in dark. After alkylation, the proteins were trapped and washed over suspension trapping columns (S-Trap micro, ProtiFi) and then digested on-column using trypsin/LysC (Promega) at 37°C overnight.

### Secretome sample preparation

Cells were seeded in 10-cm dishes and grown to confluency in DMEM supplemented with 10% FBS and antibiotics. After 2 days, cells were washed three times with 5 mL PBS to remove residual serum proteins. Serum-free DMEM was then added, and cells were incubated for 18 hours to allow protein secretion. Conditioned medium was collected and centrifuged at 1,000 × g for 3 min at 4 °C to remove cell debris, and the supernatant was retained. The medium was concentrated using Amicon Ultra centrifugal filters (3 kDa MWCO, 15 mL capacity) at 3,500 × g. Buffer exchange into 1X TBS (pH 7.6) was performed by adding 10 mL TBS and centrifuging at 3,500 × g, repeated three times, with the final volume concentrated to 300 µL. The concentrated secreted proteins were subsequently used for protein digestion and LC-MS/MS analysis, as described above, with addition of 50 mM TEAB and SDS to utilize S-trap columns for overnight protein digest with trypsin/LysC after protein reduction and alkylation.

### Liquid chromatography tandem mass spectrometry (LC-MS/MS) proteomics analysis

Digested peptides were separated on an Acclaim PepMap RSLC C18 column (75 µm x 15 cm) and eluted into the nano-electrospray ion source of an Orbitrap Eclipse™ Tribrid™ mass spectrometer (Thermo Fisher Scientific) at a flow rate of 200 nL/min. The elution gradient for the secretome samples consists of 1-40% acetonitrile in 0.1% formic acid over 220 minutes followed by 10 minutes of 80% acetonitrile in 0.1% formic acid. The gradient for all other samples consists of 1-40% acetonitrile in 0.1% formic acid over 140 minutes followed by 10 minutes of 80% acetonitrile in 0.1% formic acid. The spray voltage was set to 2.2 kV and the temperature of the heated capillary was set to 275 °C. Full MS scans were acquired from m/z 300 to 2000 at 60k resolution, and MS/MS scans following collision-induced dissociation (CID) with the collision energy at 38% were collected in the ion trap. The spectra were analyzed using SEQUEST (Proteome Discoverer 2.5, Thermo Fisher Scientific) with mass tolerance set as 20 ppm for precursors and 0.5 Da for fragments. The search output was filtered within the program to reach a 1% false discovery rate at protein level and 10% at peptide level. The quantitation was performed based on spectral counts. For secretome statistical analysis, peptide spectral counts (PSM) were normalized to Normalized Spectral Abundance Factor (NSAF) and protein identifications were required to be detected above a threshold of 30 total PSMs. Perseus v2.1.3 (Max-Planck-Institute of Biochemistry) was then used to finalize data pre-processing and statistical analyses via log2 conversion of raw NSAFs with a requirement to be detected in more than one biological replicate of each genotype. Proteins not found in an individual replicate were assigned a value of 0. GraphPad Prism (v10.8) was then used for data visualization of differential protein abundance analysis.

### Immunofluorescence

Cells were grown on glass coverslips, washed with PBS, fixed with 4% paraformaldehyde, permeabilized with 1% Triton X-100, and blocked with 2% BSA in PBST for 1 hour at room temperature. Primary antibodies were applied in 2% BSA/PBST overnight at 4°C, followed by three PBS washes (10 min each) and incubation with fluorescent secondary antibodies for 1 h at room temperature. Coverslips were washed three times with PBS, mounted using ProLong Gold antifade reagent containing DAPI (Thermo Scientific, #P36941), and sealed with clear nail polish. Imaging was performed on an Olympus IX83 Confocal Microscope.

### Micromass differentiation of growth plate-like chondroprogenitors

Immature growth plate-like chondroprogenitors (GPLCs) were maintained in DMEM/F12 (Gibco) with 10% FBS (Gibco), 100 U/mL penicillin, and 100 mg/mL streptomycin sulfate (Life Technologies), at 37°C with 5% CO2. For chondrogenesis assays, micromass cultures were established as previously described^69^ with minor modifications. Briefly, 2 x 10^5^ cells were seeded as a small droplet (10-30 µL) in the center of each well of a 24-well plate and incubated at 37 °C with 5% CO2 for 2 hours to facilitate cell attachment. Complete growth medium (500 µL) was then gently added to the side of each well without disturbing the pellet, and cultures were incubated overnight. After 24 hours, the medium was replaced with complete media + 1X insulin-transferrin-selenium (ITS, Thermo Fisher) and replenished every 3 days. Chondrogenesis was evaluated on Day 4 and Day 7 using Alcian Blue staining and gene expression was assessed at Day 7 via quantitative PCR, as described above.

For Alcian Blue staining, the culture media was aspirated, and micromass cultures were washed twice gently with PBS prior to fixation in 4% formaldehyde in PBS for 20 minutes at room temperature. Fixed micromass ) andcultures were then washed twice with PBS, briefly rinsed with deionized water (3 minutes) and stained with 1% (w/v) Alcian Blue stain (Sigma-Aldrich, #TMS-010-C) for 30 minutes at room temperature, protected from light. Excess dye was aspirated and cells were rinsed extensively using deionized water. For quantification, the dye was extracted overnight at room temperature with 6 M guanidine hydrochloride, and the absorbance was measured at 595 nm using a plate reader.

### Statistics and Reproducibility

Statistical tests, sample sizes, and numbers of biological replicates are provided in figure legends. ****p < 0.0001; ***p < 0.001; **p < 0.01; *p < 0.05. All tests were two-sided and were performed using Prism v10 (GraphPad). Error bars are shown to represent standard deviation from the mean. Each measurement was taken from a distinct biological sample.

### Data availability

Raw data for LC-MS analysis of GAGs are available at GlycoPOST^70^ under project ID GPST000656. All mass spectrometry data has been deposited to ProteomeXchange Consortium via the MassIVE partner repository (https://massive.ucsd.edu/ProteoSAFe/static/massive.jsp) with the dataset identifier MSV000099037. Raw sequencing reads and results of the RNA sequencing analysis will be made available at NCBI Gene Expression Omnibus upon publication. Any data supporting the analyses in the manuscript are available from the corresponding author upon reasonable request.

## Supplementary Information

Supplementary tables, figures, and details of experimental procedures are provided in the Supporting Information.

## Acknowledgements

We thank IBEX Technologies for their in-kind donation of heparin lyase enzymes. We thank Dr. Vicki Rosen (Harvard) for the gift of the GPLC cell line, and Dr. Richard Cummings for the gift of the *COSMC-CDG* patient fibroblast line (A20D-COSMC, M2). Recombinant ST6GALNAC1 and neuraminidase were kindly provided by Dr. Kelley Moremen (UGA). We thank Dr. Hans Wandall (University of Copenhagen) for helpful discussions regarding *O*-GalNac glycosylation and proteomics experiments. We also thank the GlycoAnalytics Core Facility at University of California, San Diego for help with analytical experiments. This work was supported by NIH grants to R.J.W. (NIAMS R21AR080957, NIGMS R35GM150736) and the NSF BioFoundry: Glycoscience Research, Education, and Training (BioF:GREAT, 240020, to L.W.). This publication includes RNA-seq data generated at the UC San Diego IGM Genomics Center utilizing an Illumina NovaSeq 6000 that was purchased with funding from a National Institutes of Health SIG grant (#S10 OD026929). Certain figure panels (Figs. 2A, 4A, 6A, 8, and Suppl.Fig. 1D, 4B) were created in BioRender.

## Author Contributions

Unless otherwise noted, X.D. performed the experimental work and analyzed the data. S.B. and P.Z. generated, processed, and analyzed the proteomics data. K.G. and K.S. generated and characterized the CD44 knockout cells and performed immunoblotting and qPCR experiments. A.B. generated, processed, and analyzed the LC-MS GAG compositional data. D.C. generated recombinant ST6GALNAC1 and sialidase for SEEL experiments. X.D., L.W., and R.J.W. wrote the paper, with input from all listed co-authors.

---

**Supplementary Figure 1.**
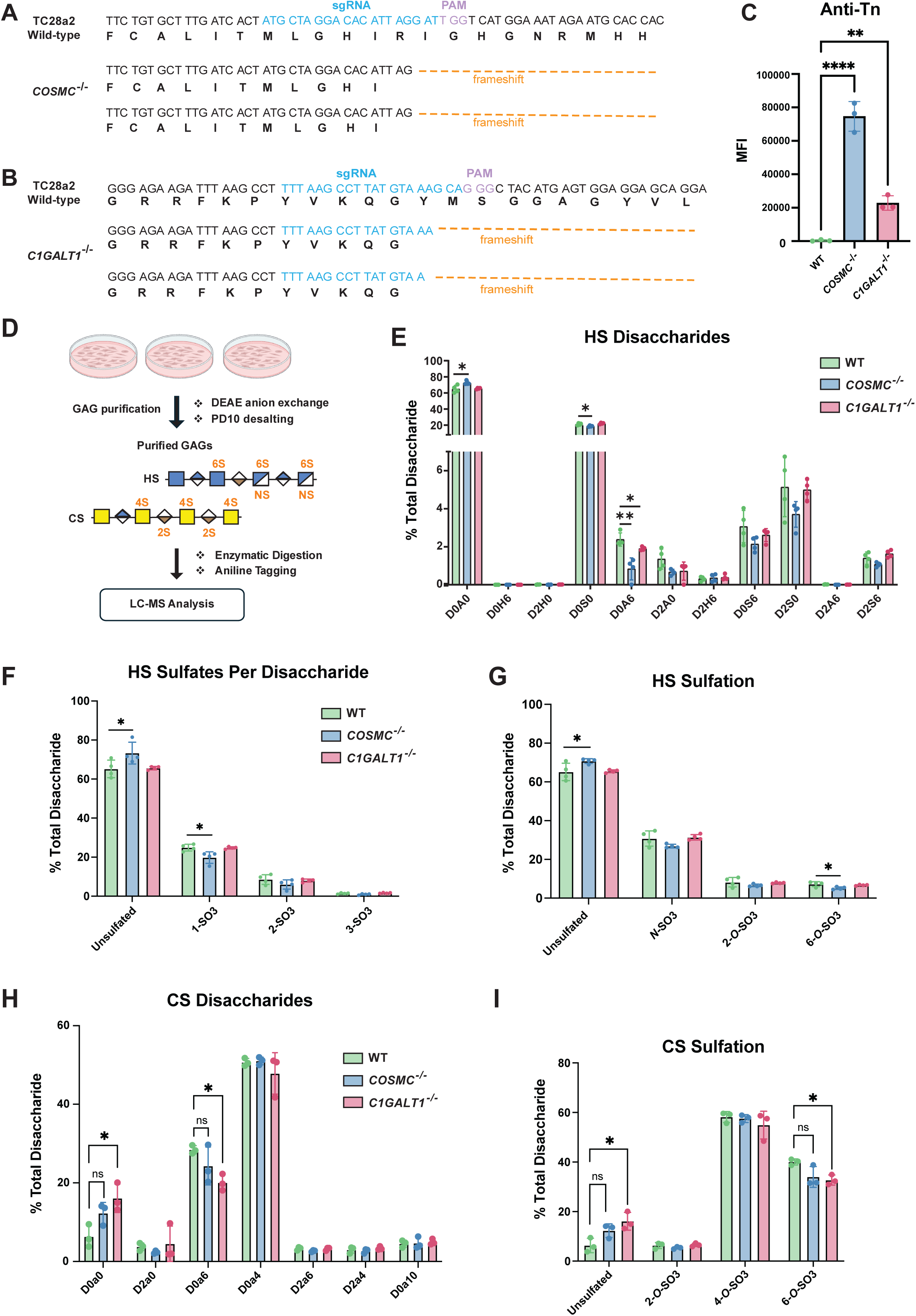
Validation and characterization of TC28a2 *COSMC^-/-^* and *C1GALT1^-/-^*knockout clones. CRISPR sgRNA targeting of human (A) *COSMC* and (B) *C1GALT1* in TC28a2 chondrocytes. Frameshift biallelic mutations in clonal cell lines for Exon 1 of *COSMC* and Exon 2 of *C1GALT1* were confirmed by Sanger sequencing. (C) Flow cytometry analysis of anti-Tn antibody binding to wild-type and *COSMC/C1GALT1* knockout cells (n = 3 independent experiments). (D) Workflow for isolation, purification, enzymatic digestion, aniline tagging, and LC-MS analysis of glycosaminoglycans. GRIL-LC-MS analysis of (E) HS disaccharides, (F) HS sulfates per disaccharide, and (G) HS sulfation in wild-type, *COSMC^-/-^*, and *C1GALT1^-/-^* knockout lines (n = 4 independent experiments). GRIL-LC-MS analysis of (H) CS/DS disaccharides and (I) CS/DS sulfation of wild-type and knockout cells (n ≥ 3 independent experiments). Data points are shown as mean ± SD; p-values were determined by student t-test with **** p < 0.0001, *** p < 0.001, ** p < 0.01, * p < 0.05.

**Supplementary Figure 2.**
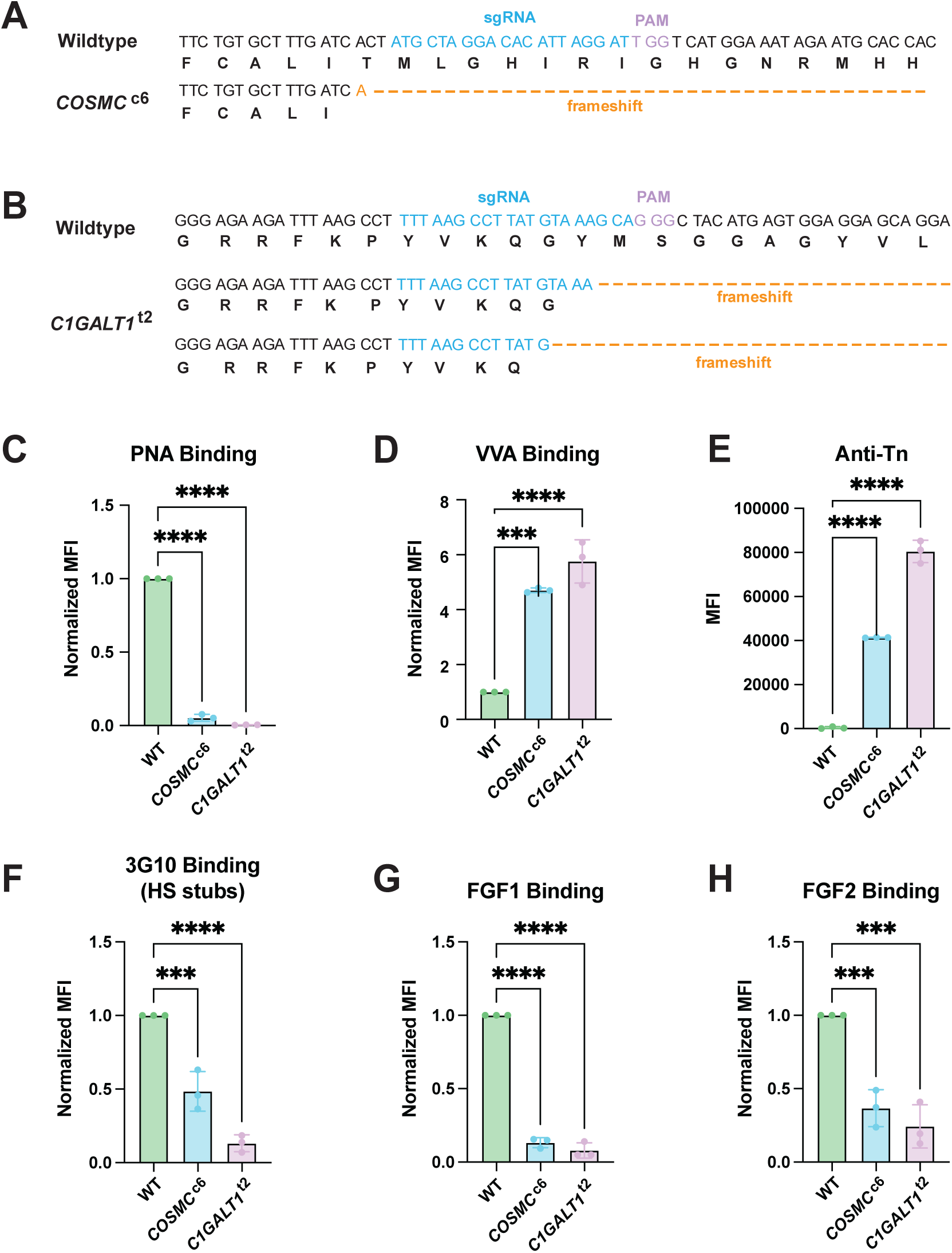
Validation and characterization of additional TC28a2 *COSMC^c6^* and *C1GALT1^t2^* knockout clones. Validation of CRISPR targeting for additional (A) *COSMC* (COSMC^c6^) and (B) *C1GALT1* (C1GALT1^t2^) knockout clones. Frameshift biallelic mutations were confirmed by Sanger sequencing. Flow cytometry analysis of (C) PNA lectin, (D) VVA lectin, (E) anti-Tn antibody, (F) 3G10, (G) FGF1, and (H) FGF2 binding in COSMC^c6^and C1GALT1^t2^ knockout clones compared with wild-type control cells. Data points are shown as mean ± SD (n ≥ 3independent experiments); p-values were determined by one-way ANOVA with Tukey post-test with \*\*\*\**p* < 0.0001, *** p < 0.001, ** p < 0.01, * p < 0.05.

**Supplementary Figure 3.**
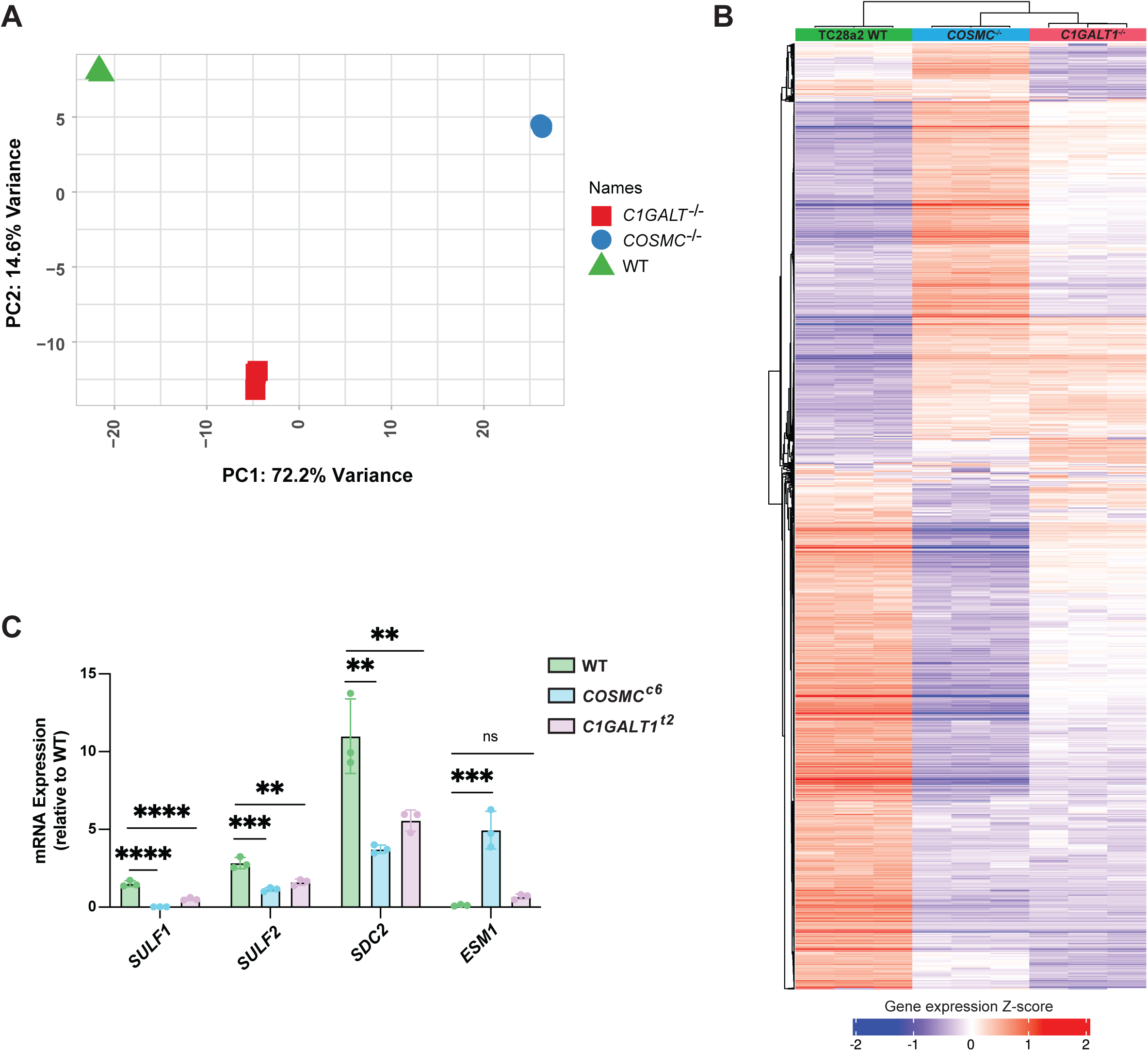
RNA-seq data analysis and qPCR validation experiments. (A) PCA analysis reveals distinct clustering of TC28a2 wild-type, *COSMC^-/-^*, and *C1GALT1^-/-^* triplicate samples. (B) Hierarchical clustering and differential expression analysis of triplicate RNA-seq datasets. (C) qPCR validation of gene expression changes in additional COSMC^c6^ and C1GALT1^t2^ knockout clones. Data points are shown as mean ± SD (n = 3 independent experiments); p-values were determined by one-way ANOVA with Tukey post-test with **** p < 0.0001, *** p < 0.001, ** p < 0.01, * p < 0.05.

**Supplementary Figure 4.**
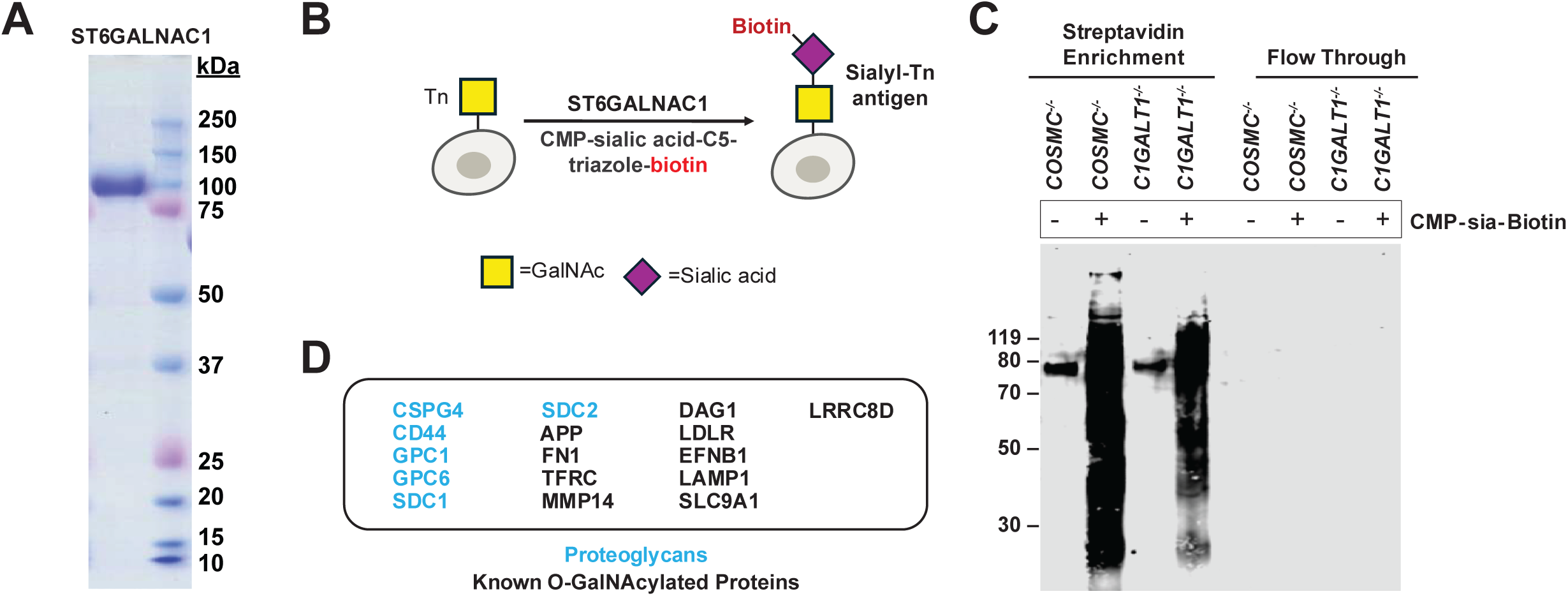
**One-step Selective exoenzymatic labeling (SEEL) with ST6GALNAC1**. (A) Commasie stained SDS-PAGE gel for purified recombinant human ST6GALNAC1-GFP fusion protein. (B) Schematic of one-step SEEL labeling of TC28a2 cells with ST6GALNAC1 and CMP-Neu5Ac-C5-triazole-biotin. (C) Representative western blot showing streptavidin enrichment versus flow through from SEEL labeling of *COSMC^-/-^* and *C1GALT1^-/-^*cells. The one-step SEEL reactions with ST6GALNAC1 were performed as described in the Methods section. (D) The SEEL labeling method detected a number of known *O*-GalNAcylated glycoproteins (black), including multiple heparan sulfate proteoglycans (blue).

**Supplementary Figure 5.**
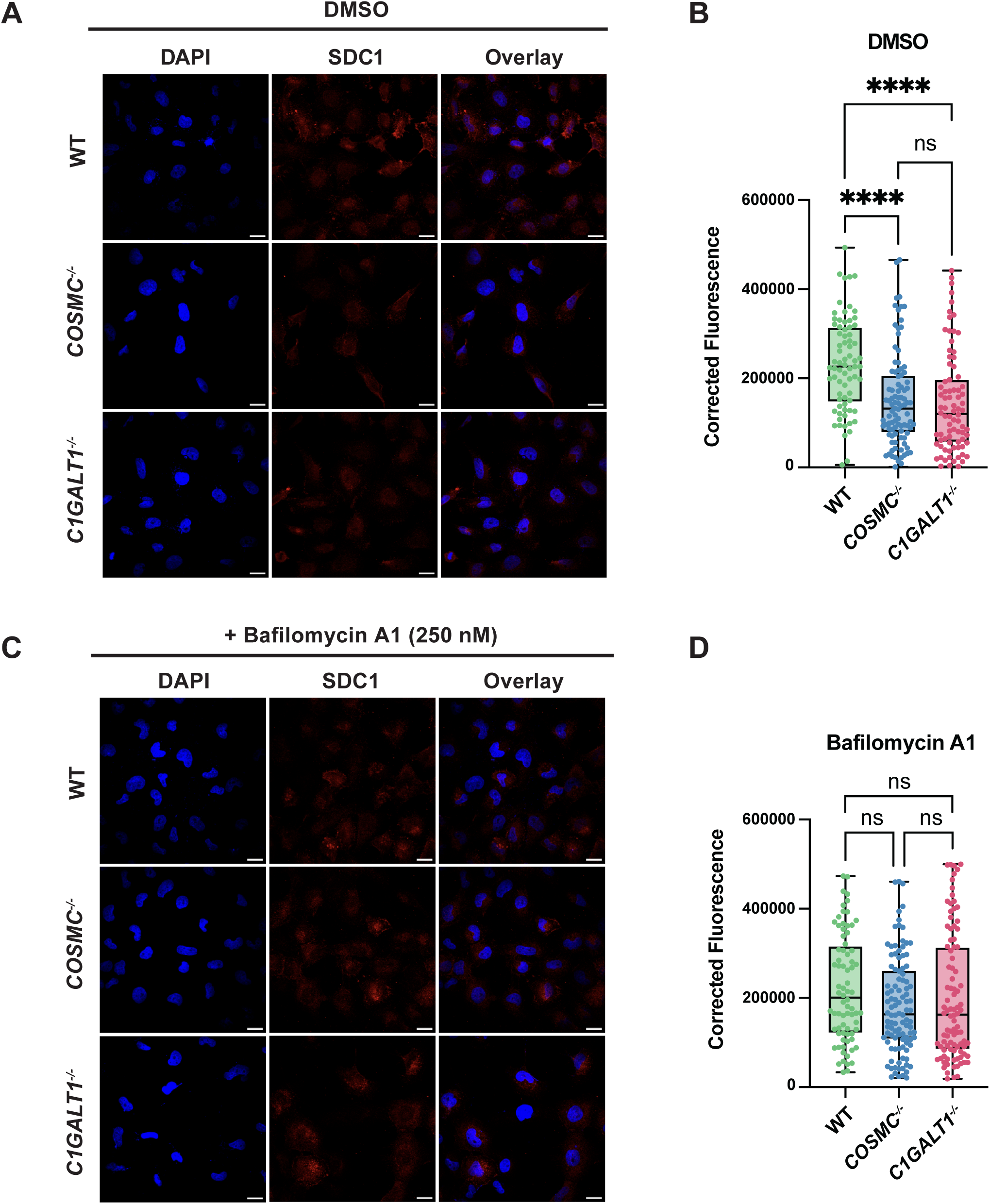
Immunofluorescence images of SDC1 staining in TC28a2 chondrocytes. (A) Representative confocal images of SDC1 (red) and nuclei (DAPI; blue) in DMSO-treated cells. The scale bar represents 20 nm. (B) Quantification of SDC1 fluorescence intensity using ImageJ. (C) Representative confocal images of SDC1 (red) and nuclei (DAPI; blue) in cells after treatment with bafilomycin A1 (250 nM) for 16 hours. The scale bar represents 20 nm. (B) Quantification of SDC1 fluorescence intensity in bafilomycin A1-treated cells versus wild-type controls. Data points are shown as mean ± SD (n = 25-35 cells from three independent biological replicates); p-values were determined by one-way ANOVA with Tukey post-test with **** p < 0.0001.

**Supplementary Figure 6.**
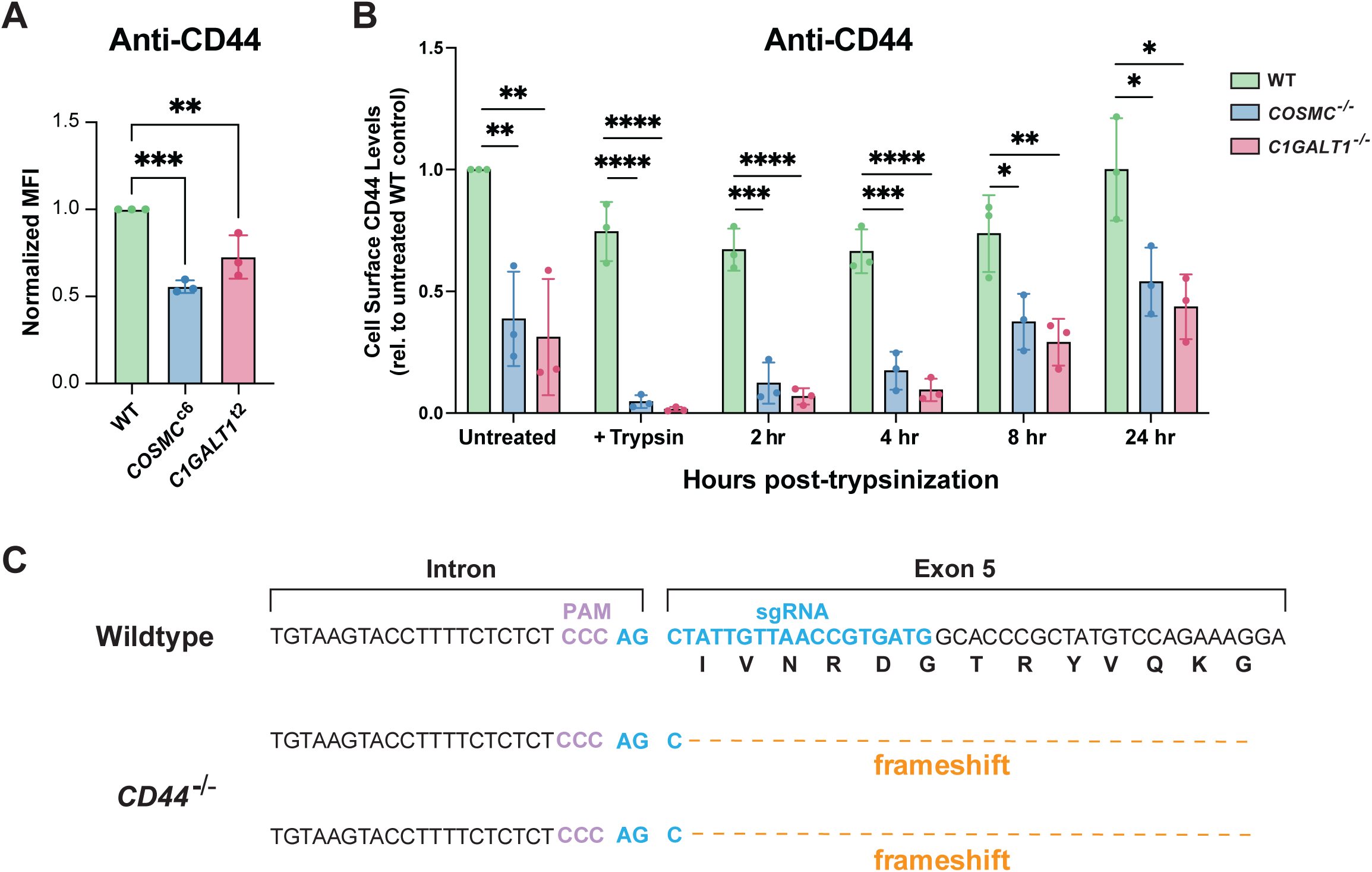
Anti-CD44 binding assays and *CD44^-/-^* knockout clone validation. (A) Flow cytometry analysis of cell surface CD44 levels in *COSMC^c6^* and *C1GALT1^t2^*knockout clones. (B) Time course quantification of cell surface CD44 levels via flow cytometry prior to and after trypsinization (0-24 hours). Data is normalized to untreated wild-type cells. Data points are shown as mean ± SD (n = 3 independent experiments); p-values were determined by one-way ANOVA with Tukey post-test with **** p < 0.0001, *** p < 0.001, ** p < 0.01, * p < 0.05. (C) CRISPR sgRNA targeting of human *CD44* in TC28a2 chondrocytes. (C) Frameshift biallelic mutations in a clonal cell line for Exon 5 of *CD44* were confirmed by Sanger sequencing.

**Supplementary Figure 7.**
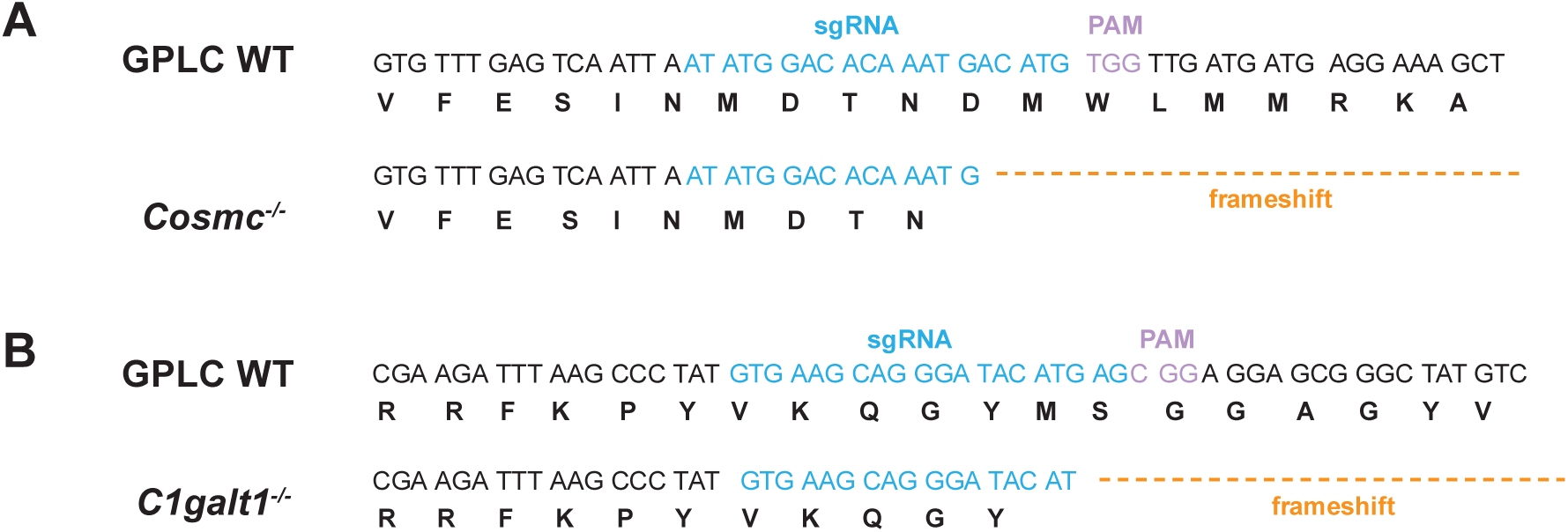
Validation of murine GPLC *Cosmc^-/-^* and *C1galt1^-/-^*knockout clones. CRISPR sgRNA targeting of mouse (A) *Cosmc* and (B) *C1galt1* in murine growth plate- like chondroprogenitors (GPLCs). Frameshift biallelic mutations in a clonal cell line for Exon 1 of *Cosmc* and Exon 2 of *C1galt1* were confirmed by Sanger sequencing.

