## Supplementary Information for "Mucin-type *O*-glycans regulate proteoglycan stability and chondrocyte maturation"

**This PDF file includes:**

Supplementary Tables 1-9
Supplementary Figures 8-9

**Supplementary Table 1:**
sgRNA sequences used for CRISPR/Cas9 gene editing

| Gene | Species | Guide Sequence (5'→3') |
|---|---|---|
| Non-targeting | – | ACGTTCGAGTACGACCAGCT |
| *COSMC (C1GALT1C1)* | Human | ATGCTAGGACACATTAGGAT |
| *C1GALT1* | Human | TTTAAGCCTTATGTAAAGCA |
| *Cosmc (C1galt1c1)* | Mouse | ATATGGACACAAATGACATG |
| *C1galt1* | Mouse | GTGAAGCAGGGATACATGAG |
| *CD44* | Human | CATCACGGTTAACAATAGCT |

**Supplementary Table 2:**
PCR primer sequences used for genotyping CRISPR/Cas9 engineered cells

| Gene | Forward Primer (5'→3') | Reverse Primer (3'→5') |
|---|---|---|
| *COSMC* | TGATTTCAAGCTTGGGAACCTTT | CATCCCTCCCTGTTCAGGAC |
| *C1GALT1* | CCCTGCTGTGGGACTGAAAA | GGTGTTCTGGCACAAAGGGA |
| *CD44* | ACATAGACGTTGCTGAGAACC | TTCCATGCACCTTCCCAGAG |
| *Cosmc (m)* | AGGGAAACAAGTGGGAGAATGA | TCCCTGAATGGCATCACCAG |
| *C1galt1 (m)* | ACCATCTGCAAGCCCCTAA | TACTCCGGCGTATTTCAGGC |

**Supplementary Table 3:**
Primer sequences used for quantitative PCR analysis of gene expression

| Gene | Forward Primer Sequence (5'→3') | Reverse Primer Sequence (3'→5') |
|---|---|---|
| *YWHAZ* | CCTGCATGAAGTCTGTAACTGAG | GACCTACGGGCTCCTACAACA |
| *SDC2* | GCTGTTGGTGTATCGCATGA | ACTGGATGGTTTGCGTTCTC |
| *SULF1* | GAAGGAGAAGAGACGGCAGA | CAGAAAGATCCCAGGTTCCA |
| *SULF2* | CACTGGCAAGTACGTCCACAA | CTATTGAGGTACACCCCAAAGG |
| *ESM1* | GAGAAACTTGCTACCGCACA | CCCATTAGAAGGCTGACACC |
| *Gapdh (m)* | GGTGCTGAGTATGTCGTGGA | CCTTCCACAATGCCAAAGTT |
| *Sox9 (m)* | AGTACCCGCATCTGCACAAC | TACTTGTAATCGGGGTGGTCTTTC |
| *Col2a1 (m)* | AGGGCAACAGCAGGTTCACATAC | TGTCCACACCAAATTCCTGTTCA |
| *Acan (m)* | GCTGCAGTGATCTCAGAAGAAG | GATGGTGAGGGAAGACCCTA |

**Supplementary Table 4:**
HS disaccharide composition for TC28a2 WT, *COSMC* KO, and *C1GALT1* KO cells

| Disaccharide Structure | | Abundance (% Total Disaccharide)[c] | | |
|---|---|---|---|---|
| Structure Code[a] | Unit Formula[b] | WT (% Total HS) | *COSMC*$^{-/-}$ (% Total HS) | *C1GALT1*$^{-/-}$ (% Total HS) |
| D0A0 | ΔUA-GlcNAc | 64.97 ± 4.50 | 72.48 ± 2.34 | 65.50 ± 0.83 |
| D0H6 | ΔUA-GlcNH$_2$6S | - | - | - |
| D2H0 | ΔUA2S-GlcNH$_2$ | - | - | - |
| D0S0 | ΔUA-GlcNS | 21.13 ± 1.27 | 18.72 ± 0.81 | 22.13 ± 0.86 |
| D0A6 | ΔUA-GlcNAc6S | 2.38 ± 0.35 | 0.83 ± 0.58 | 1.90 ± 0.09 |
| D2A0 | ΔUA2S-GlcNAc | 1.35 ± 0.52 | 0.67 ± 0.15 | 0.72 ± 0.48 |
| D2H6 | ΔUA2S-GlcNH$_2$6S | 0.31 ± 0.12 | 0.36 ± 0.20 | 0.39 ± 0.14 |
| D0S6 | ΔUA-GlcNS6S | 3.06 ± 0.85 | 2.14 ± 0.38 | 2.61 ± 0.34 |
| D2S0 | ΔUA2S-GlcNS | 5.13 ± 1.55 | 3.70 ± 0.67 | 4.99 ± 0.51 |
| D2A6 | ΔUA2S-GlcNAc6S | - | - | - |
| D2S6 | ΔUA2S-GlcNS6S | 1.40 ± 0.32 | 1.07 ± 0.11 | 1.63 ± 0.19 |

[a] The disaccharide structure code is described in (Lawrence, et al. Nat. Methods 2008)
[b] ΔUA = 4,5-unsaturated uronic acid
[c] –, not detected

**Supplementary Table 5:**
HS sulfation and *N*-substitution of glucosamine units for TC28a2 WT, *COSMC* KO, and *C1GALT1* KO cells

| HS Sulfation | Constituents/100 disaccharides | | |
|---|---|---|---|
| | WT | *COSMC*$^{-/-}$ | *C1GALT1*$^{-/-}$ |
| Unsubstituted glucosamine | 65.16 ± 4.47 | 70.74 ± 1.22 | 65.48 ± 0.74 |
| *N*-sulfoglucosamine | 30.76 ± 3.94 | 26.83 ± 0.95 | 31.39 ± 1.42 |
| Uronyl-2-*O*-sulfates | 8.23 ± 2.44 | 6.58 ± 0.65 | 7.86 ± 0.31 |
| Glucosaminyl 6-*O*-sulfates | 7.24 ± 1.26 | 5.15 ± 0.64 | 6.68 ± 0.28 |

**Supplementary Table 6:**
HS sulfate groups per disaccharide of glucosamine units for TC28a2 WT, *COSMC* KO, and *C1GALT1* KO cells

| HS Sulfates/disaccharide | Abundance (% Total Disaccharide) | | |
|---|---|---|---|
| | **WT** | ***COSMC$^{-/-}$*** | ***C1GALT1$^{-/-}$*** |
| 0 $SO_3$ | 65.20 ± 4.48 | 73.29 ± 5.57 | 65.60 ± 0.85 |
| 1 $SO_3$ | 24.87 ± 1.72 | 19.78 ± 2.98 | 24.77 ± 0.59 |
| 2 $SO_3$ | 8.53 ± 2.48 | 6.01 ± 2.40 | 8.00 ± 0.79 |
| 3 $SO_3$ | 1.40 ± 0.32 | 0.92 ± 0.33 | 1.63 ± 0.19 |

**Supplementary Table 7:**
CS/DS disaccharide composition of TC28a2 WT, *COSMC* KO, and *C1GALT1* KO cells

| Disaccharide Structure | | Abundance (% Total Disaccharide) | | |
|---|---|---|---|---|
| **Structure Code[a]** | **Unit Formula[b]** | **WT (% Total CS)** | ***COSMC$^{-/-}$*** **(% Total CS)** | ***C1GALT1$^{-/-}$*** **(% Total CS)** |
| D0a0 | ΔUA-GalNAc | 6.32 ± 2.89 | 12.29 ± 2.75 | 16.44 ± 4.21 |
| D2a0 | ΔUA2S-GalNAc | 3.76 ± 0.97 | 2.35 ± 0.37 | 1.83 ± 0.15 |
| D0a6 | ΔUA-GalNAc6S | 28.44 ± 1.04 | 24.31 ± 4.84 | 20.07 ± 2.08 |
| D0a4 | ΔUA-GalNAc4S | 50.68 ± 0.99 | 51.04 ± 0.98 | 51.16 ± 0.48 |
| D2a6 | ΔUA2S-GalNAc6S | 3.28 ± 0.41 | 2.68 ± 0.21 | 3.24 ± 0.31 |
| D2a4 | ΔUA2S-GalNAc4S | 2.98 ± 0.82 | 2.69 ± 0.55 | 3.42 ± 0.42 |
| D0a10 | ΔUA-GalNAc4S6S | 4.53 ± 0.89 | 4.65 ± 1.48 | 3.83 ± 2.47 |

**Supplementary Table 8:**
CS/DS sulfation of *N*-acetyl-galactosamine units in TC28a2 WT, *COSMC* KO, and *C1GALT1* KO cells

| CS/DS Sulfation | Abundance (% Total Disaccharide) | | |
|---|---|---|---|
| | **WT** | ***COSMC$^{-/-}$*** | ***C1GALT1$^{-/-}$*** |
| Unsubstituted *N*-acetyl-galactosamine | 6.32 ± 2.89 | 12.29 ± 2.75 | 16.44 ± 4.21 |
| Uronyl-2-*O*-sulfates | 6.27 ± 1.22 | 5.36 ± 0.46 | 6.66 ± 0.56 |
| GalNAc 4-*O*-sulfates | 58.16 ± 2.20 | 57.54 ± 1.56 | 57.08 ± 1.88 |
| GalNAc 6-*O*-sulfates | 40.02 ± 1.23 | 33.98 ± 4.19 | 28.99 ± 4.57 |

**Supplementary Table 9:**
CS/DS sulfate groups per disaccharide of *N*-acetyl-galactosamine units in Tc28a2 WT, *COSMC* KO, and *C1GALT1* KO cells

| CS/DS Sulfation | Abundance (% Total Disaccharide)[a] | | |
|---|---|---|---|
| | WT | *COSMC*$^{-/-}$ | *C1GALT1*$^{-/-}$ |
| 0 $SO_3$ | 6.32 ± 2.89 | 12.29 ± 2.75 | 16.44 ± 4.21 |
| 1 $SO_3$ | 82.88 ± 1.56 | 77.70 ± 4.38 | 73.07 ± 1.69 |
| 2 $SO_3$ | 10.81 ± 2.11 | 10.01 ± 1.91 | 10.49 ± 2.77 |
| 3 $SO_3$ | - | - | - |

[a] –, not detected

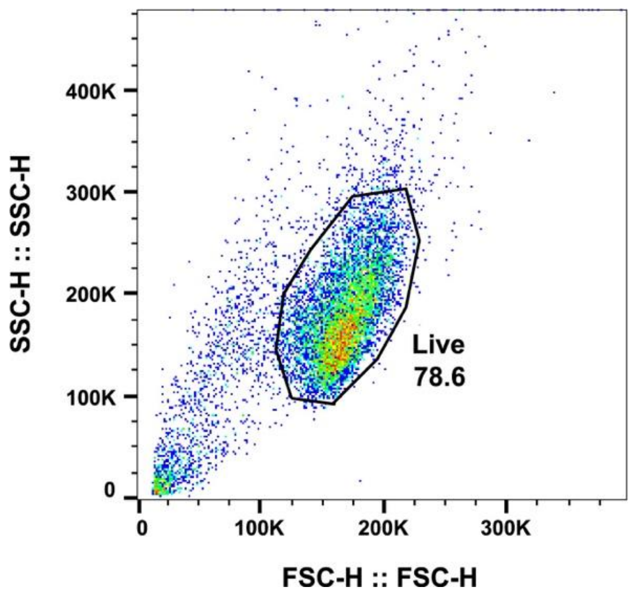


**Supplementary Figure 8. General flow cytometry gating strategy.** Cells were gated based on forward and side scattering for analysis of flow cytometry data.

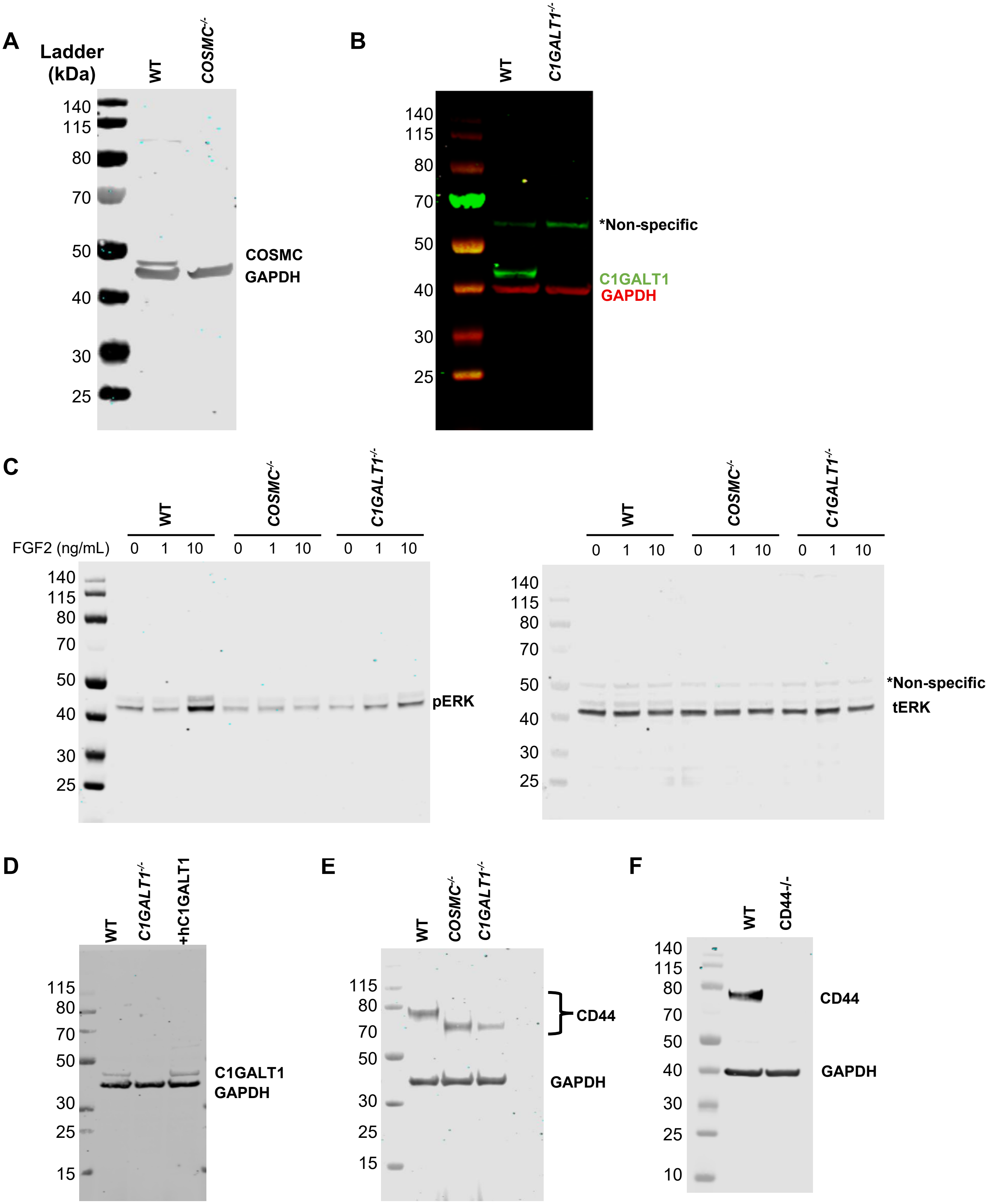


**Supplementary Figure 9. Source Data.** Uncropped western blot source images for (A) Figure 1B, (B) Figure 1C, (C) Figure 2E, (D) Figure 2G, (E) Figure 6B, and (F) Figure 6F.
